# High-resolution taxonomic profiling and metatranscriptomics identify microbial, biochemical, host and ecological factors in peri-implant disease

**DOI:** 10.1101/2025.06.23.661096

**Authors:** Szymon P. Szafrański, Amruta A. Joshi, Matthias Steglich, Ines Yang, Taoran Qu, Wiebke Behrens, Uthayakumar Muthukumarasamy, Damianos Melidis, Paula Schaefer-Dreyer, Jasmin Grischke, Jan Hegermann, Wolfgang Nejdl, Susanne Häussler, Meike Stiesch

## Abstract

Biofilm-associated diseases like peri-implant mucositis (PIM) and peri-implantitis (PI) are significant clinical challenges affecting millions of dental implant patients globally. Although studies have described the role of microbial, host, or environmental factors in disease development, their complex interplay, particularly during dysbiosis remains poorly understood. This cross-sectional study characterized the microbiome composition and metatranscriptomes of 125 peri-implant biofilms from 48 individuals uncovering molecular signatures linked to peri-implant health (PIH), PIM, and PI. Distinct variations were observed in biofilm amount, composition, activity, phage populations and host response. Biofilms were categorized into four community types (CTs) based on the bacterial transcriptional activity: one linked to PIH, one to PI, and two to PIM. PIH and PIM were primarily characterized by aerotolerant taxa with increased anabolic processes, while PI was dominated by obligate anaerobes with complex biofilm morphology, and heightened catabolic activity and virulence. PIM samples, relative to PIH were characterized by biofilm expansion with minimal functional changes, except for the *Neisseria*-rich PIM subtype showing higher pyruvate and lipoic acid metabolism. The phagome mirrored the bacterial compositional variations across disease states. Furthermore, human transcriptome responses varied indicating increased keratinization in PIH, enhanced expression of ribosome components in PIM, and inflammatory signaling and hypoxia in PI. Additionally, we identified complex species-enzyme, phage-bacteria, and host-microbe associations within the peri-implant ecosystem. Our integrative multi-omics approach provides a comprehensive view of microbial, biochemical, host, and ecological factors associated with dysbiosis, offering novel insights into peri-implant disease dynamics.

**Importance:** Peri-implant mucositis and peri-implantitis are highly prevalent inflammatory conditions that compromise the long-term survival and success of dental implants, yet their underlying biological mechanisms are largely unresolved. While next-generation sequencing has advanced our understanding of microbial composition across health and peri-implant diseases, it falls short of capturing microbial activity and the broader molecular context of peri-implant dysbiosis. Metatranscriptomics overcomes this limitation by profiling actively transcribed genes within the biofilm, offering direct insights into microbial community functions. In this study, we integrated full-length 16S rRNA gene amplicon sequencing with metatranscriptomic profiling to simultaneously assess microbial taxonomy, functional activity, phage dynamics, and host gene expression in peri-implant biofilms. Importantly, we provide a systems-level view and report previously undescribed associations between different molecular signatures in peri-implant ecosystem.

## Introduction

Millions of patients have received dental implants as an effective solution for tooth replacement^1^. However, the growing number of dental implant placement has been a accompanied by a rise in biofilm-associated peri-implant diseases. Peri-implant mucositis (PIM), a reversible infection of the peri-implant mucosa, has a prevalence of 43%, while peri-implantitis (PI), which involves progressive loss of the supporting bone, affects more than 25% of implants within five years of placement^2, 3^. These peri-implant diseases impact patients’ quality of life and pose a substantial public health burden^4^. The primary etiological factor for both PIM and PI is an imbalance within the microbial community characterized by uncontrolled biofilm expansion and the emergence of more virulent species^5–14^. This dysbiosis triggers a host immune response which, if unsuccessful in clearing the infection, becomes dysregulated^8, 15, 16^. The resulting immune dysfunction leads to tissue destruction, creating an environment that further promotes dysbiosis. Traditionally, peri-implant disease has been considered closely related to periodontal disease, however, emerging evidence suggests that the two ecosystems differ significantly^5, 8, 14, 17^. While inter-subject variability in the submucosal microbiome has been observed among individuals with peri-implant disease, it remains unclear whether distinct community types, similar to those identified in periodontal disease, exist^18, 19^.

Despite advances in molecular methods, the mechanisms underlying the development of the complex, multifactorial peri-implant conditions remain to be elucidated^8, 20^. Metataxonomics and metagenomics overcame the limitations of early culture-based and DNA-hybridization methods, and enabled comprehensive profiling of the peri-implant microbiome and its functional potential^7, 11, 12, 14^. However, short-read 16S sequencing lack high resolution of taxonomic diversity (except metagenomics) and falls short in providing direct insights into bacterial activities, which can be uncovered by metatranscriptomics (MTX). Additionally, MTX provides a unique perspective on interspecies interactions, enabling predictions about their role in ecosystem dynamics and the development of dysbiosis^21, 22^. Interestingly, MTX sequences from peri-implant biofilms also contain host and bacteriophage RNA offering important insights into host-biofilm and phage-bacteria interactions^23, 24^. However, technical challenges have limited the application of MTX on peri-implant biofilms, confining previous studies to two small patient cohorts^5, 16^.

A multi-omics approach, integrating full-length 16S rRNA gene amplicon sequencing (full-16S) and MTX, represents a promising research direction for characterizing biofilm interactions and revealing the underlying mechanisms of microbiome dynamics and disease development^25–28^, especially considering the complex and multi-dimensional nature of the submucosal ecosystem^22^. In this study, we employed paired full-16S and MTX in the largest cohort of dental implant patients to date, aiming to advance a systems-level understanding of the peri-implant disease etiology. This integrative approach allowed a comprehensive characterization of the peri-implant ecosystem across three diagnosis groups, including the compositions and activity of bacterial species and their community types, bacteriophages, enzymes, host genes, as well as key interactions among these components.

## Results

In this study we analyzed the peri-implant microbiota composition and activity, including bacteriophages, and patient transcriptional responses from 125 implants in 48 patients (**Fig. 1a**). Implants were categorized based on their peri-implant disease status according to the ‘Classification of Periodontal and Peri-Implant Diseases and Conditions’^29^. 56 implants were diagnosed as healthy (PIH), 37 with peri-implant mucositis (PIM) and 32 with peri-implantitis (PI). PI-associated implants showed significantly higher plaque index scores, poor oral hygiene, probing depths, suppuration and history of periodontal disease (**Supplemental Table 1**). PI was characterized by a higher incidence of pus formation and pain. We obtained paired full-length 16S rRNA gene amplicon profiles (full-16S) and metatranscriptomes (MTX) for 125 implants. We performed full-16S instead of shotgun metagenomics. While shotgun approaches can complement MTX as a mapping reference, they are less feasible here due to the extremely low biomass of peri-implant biofilm samples, especially from healthy sites. We used an RNA-focused co-isolation protocol to preserve RNA integrity, which limits DNA quality. Full-16S allowed high-resolution taxonomic profiling from the same RNA-coisolated samples used for MTX, helping to minimize batch effects from separate sampling. In this setting, it provided a practical and informative alternative. Multiple samples per patient captured variability. Profiles were averaged by diagnosis to control for patient effects. MTX data concurrently assessed bacteria (MTX-B), phages (MTX-P), and host response (MTX-H). Comprehensive details on sequencing results are available in **Supplemental Table 2**.

**Figure 1.**
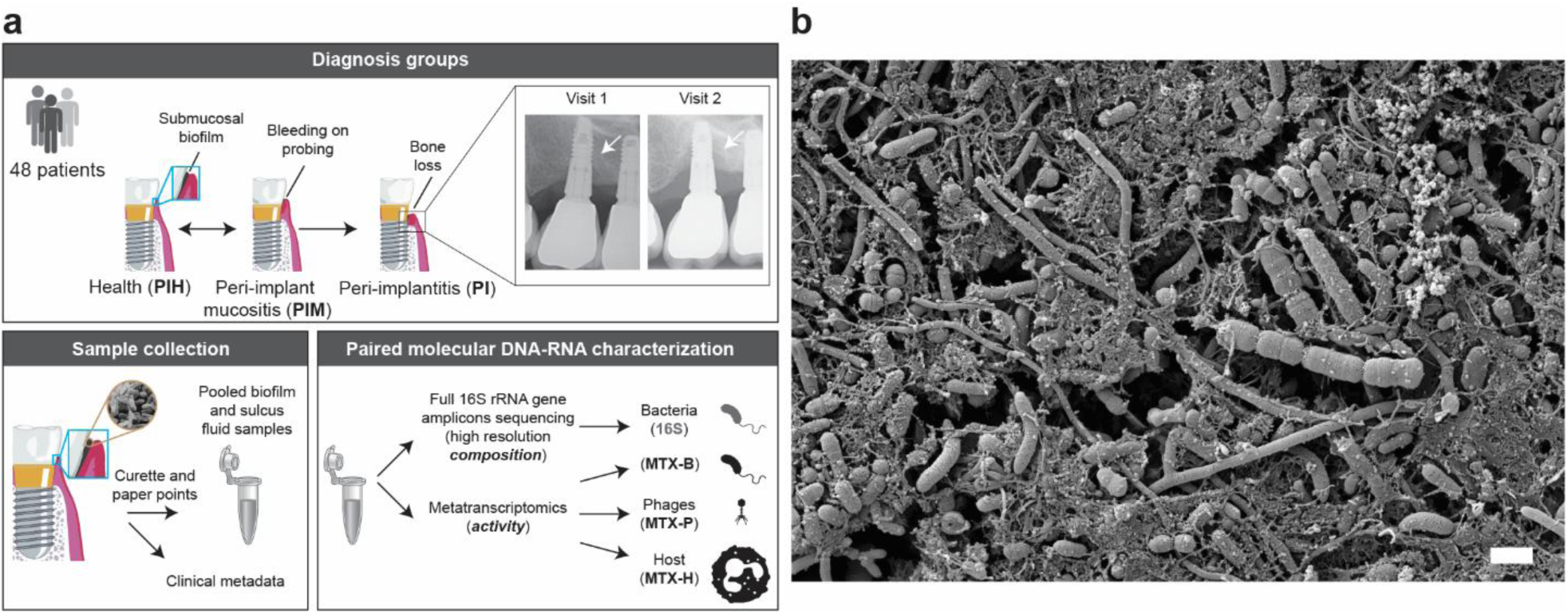
Study design. **(A)** Peri-implant diseases (top). Bone loss is an indicator of peri-implantitis. Radiographs showing crestal bone level in a representative clinical case with developing peri-implant disease. Collection of clinical samples (bottom left). Characterization of clinical samples (bottom right): For each implant, DNA- and RNA-based analyses were perfomed on the different nucleotide fractions of each sample. B-MTX, P-MTX and H-MTX analyses were performed on the same sulcus fluid samples. (**B)** Scanning electron micrograph of a submucosal implant-associated biofilm scraped with a curette from an implant diagnosed with peri-implantitis. Length of scale bar: 1µm. Implant image was modified from Zheng, H. et al. Sci Rep. 5, 2015.

### Peri-implantitis exhibits dominance of anaerobic taxa, including many unnamed fastidious species

To identify the relative abundances of taxa that are differentially associated with PIH, PIM and PI, we analyzed the composition of peri-implant communities (**Supplemental Table 3**). Major compositional differences occurred at the class level between disease states (**Supplemental Fig. 1**, PERMANOVA pseudo F =3.15, *P*=0.0028). PIH and PIM were generally characterized by a high abundance of aerobic or aerotolerant microorganisms of Bacilli, Actinobacteria, and Gammaproteobacteria while PI showed dominance of strict anaerobes represented by Bacteroidia, Clostridia, and Deltaproteobacteria (Kruskal Wallis adj. *P* ≤ 0.05). The biofilm composition in PIM was intermediate between that of PIH and PI, more closely resembling PIH (PERMANOVA *P* = 0.65 versus *P* = 0.01 for the PIM-PI comparison).

Abundances of PI-associated classes positively correlated with clinical indicators of disease severity (**Supplemental Fig. S1c** and **d**). Few outlier samples with taxonomic profiles not typical for a given diagnosis group were observed.

At lower taxonomic levels, we identified a total of 99 different genera and 756 species-level phylotypes (**Fig. 2**, **Supplemental Table 4, 5**). Microbial diversity, as reflected by the Shannon diversity index (H’) was higher in PI compared to PIH (3.4 ± 0.5 vs 3.1 ± 0.5, mean ± SD, *P* = 0.02). In total, 23 genera and 28 species were dominant (average relative abundance of ≥ 1% in at least one diagnosis group), 30 genera and 34 species were core (present in ≥ 75% of samples in at least one group), and 13 genera and 63 species were associated with different diagnosis groups (FDR-corrected *P* < 0.05, **Supplemental Table 4, 5**). At the genus level, *Streptococcus* was dominant and core in both PIH and PIM. PIM showed significantly higher abundances of *Shuttleworthia*, while *Aggregatibacter* and *Peptostreptococcus* were less abundant compared to PIH. PI showed expansion of *Dialister*, *Mogibacterium*, *Prevotella*, *Parvimonas*, *Pseudoramibacter*, *Eubacterium* outgrowing *Streptococcus* and 4 other genera when compared to PIH and PIM. At the species level, abundant species formed three major clusters of samples, each primarily associated with one diagnosis group (**Fig. 2**). In our study, we identified 228 of the 325 unnamed provisional species from the oral cavity^30^, assigned with Human Microbiome Taxon (HMT) numbers. Of these, 157 met the relative abundance threshold of 0.1% across all samples (**Supplemental Table 5**). These species corresponded to a mean of 9.3% of the analyzed communities (maximum 36%) with *Fusobacterium* sp. HMT-203 being the most abundant. Unnamed species with HMT numbers were more abundant in PIM (10.6 ± 7.7, mean ± sd, *P* = 0.013) and PI (12.1 ± 6.8, mean ± sd, *P* = 0.0004) compared to PIH (7.0 ± 5.9). In summary, microbial communities on implants exhibited a remarkable complexity, comprising a substantial proportion of understudied and yet unnamed species. Notably, overall microbial diversity and abundance of anaerobic taxa increased in PI.

**Figure 2.**
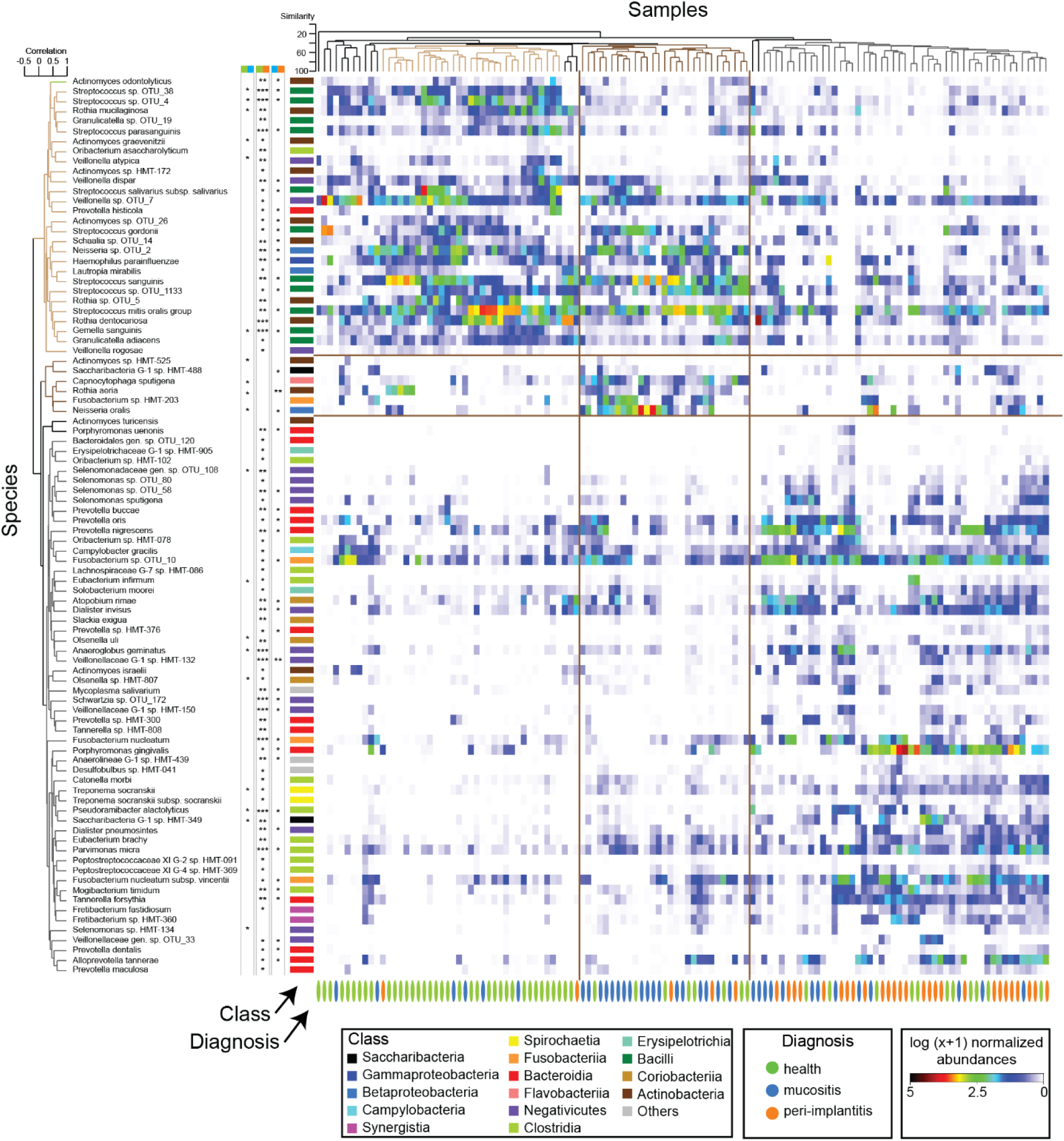
High-resolution composition of implant-associated biofilms. Heatmap indicating relative abundances of the top species-level taxa selected by CAP analysis (Pearson |r| > 0.4) across diagnosis groups. Horizontal coloured bars indicating bacterial class, vertical bars below heatmap indicating the diagnosis associated with the sample. Samples ordered by hierarchical clustering of Bray–Curtis similarity values calculated for log (x +1) transformed standardized species abundance profiles. Species ordered by hierarchical clustering of Spearman rank correlation values calculated for log (x +1) transformed standardized species abundance profiles. Significant *P-*values for Mann-Whitney U test for pair-wise comparisons and Benjamini Hochberg correction across diagnosis groups. *** *P* ≤ 0.0001; ** *P* ≤ 0.001; * *P* ≤ 0.05

### Peri-implantitis is characterized by high bacterial RNA yields and complex biofilm morphology

To capture microbial community functions beyond the microbiota composition detailed above, we incorporated a metatranscriptomic perspective in our investigation. The amount of bacterial RNA isolated from peri-implant samples and reflecting absolute levels of biofilm activity, was determined by multiplying the total RNA by the proportion of reads mapped to bacteria. The results were validated by the rRNA signals observed in Bioanalyzer profiles of the total RNA. The total amount of bacterial RNA varied significantly across diagnosis groups (Kruskal-Wallis, *H* = 24.5, *P* = 4.7 · 10^-6^). The median bacterial RNA amount increased from 1 ng in PIH to 10 ng in PIM (post-hoc Dunn’s test, Bonferroni-adjusted *P* = 0.0096), and 50 ng in PI (*P* = 2.1 · 10^-6^ and *P* = 0.04 compared to PIH and PIM, respectively). A striking difference of four orders of magnitude was observed in bacterial RNA levels across samples, ranging from 0.05 ng to 1,500 ng. Notably, samples with exceptionally high RNA yields corresponded to high amounts of plaque observed macroscopically during sampling and processing. Electron microscopy was performed on four PI samples with extensive biofilm colonization to characterize their morphology (**Fig. 1b**, **Supplemental Fig. 2**). These biofilms displayed complex structures which included cell filaments, bacterial aggregates, extracellular networks and localized granules of extracellular polymeric substances (EPS), as well as features involved in biofilm stability, communication and dispersal/remodelling, such as vesicles and motile flagellated cells.

### Metatranscriptomics revealed four major active community types (CT) in peri-implant biofilms

Metatranscriptomics allowed us to capture significant differences in activity distribution by analyzing the overall mRNA abundances across bacterial classes (**Fig. 3**, **Supplemental Fig. 3**). Based on their activity profiles, biofilms at class level showed statistically significant clustering (type 1 SIMPROF test, significance level of 0.1%). Importantly, clustering was not based on arbitrary similarity thresholds but was determined using the SIMPROF (similarity profile) algorithm, which employs permutation-based testing to identify statistically significant groupings within hierarchical clustering, thereby offering a robust approach to identify genuine clusters. Nine clusters of active communities, with four major clusters (community types CTs I-IV, **Fig. 3a**) covering most samples, were identified. Each CT displayed a unique profile of active bacterial classes. CT I was primarily associated with PIH samples and negatively associated with clinical measures reflecting disease severity. CT II almost exclusively included PIM samples, while CT III exhibited PIM as the highest proportion. CT IV had the highest proportion of PI samples. Within some individuals, some biofilms from different implants with the same diagnosis showed distinct CTs (**Fig. 3b**). CT I was characterized by high Bacilli activity; mainly from *Streptococcus*. CT II was dominated by Betaproteobacteria activity, particularly *Neisseria*. CT III showed elevated activity of Bacteroidia (*Prevotella*) and Fusobacteriia (*Fusobacterium*), while CT IV exhibited even higher Bacteroidia activity, specifically from *Tannerella* and *Porphyromonas*, in contrast to CT III along with increased Fusobacteria activity. The species co-occurence network visualized through ordination showed unique networks of pairwise correlations between active species within CTs. CT II displayed the highest number of co-occuring taxa (**Fig. 3d**). Hence, clustering of active biofilm profiles identified four major community types, with CT II and CT III being predominantly associated with PIM.

**Figure 3.**
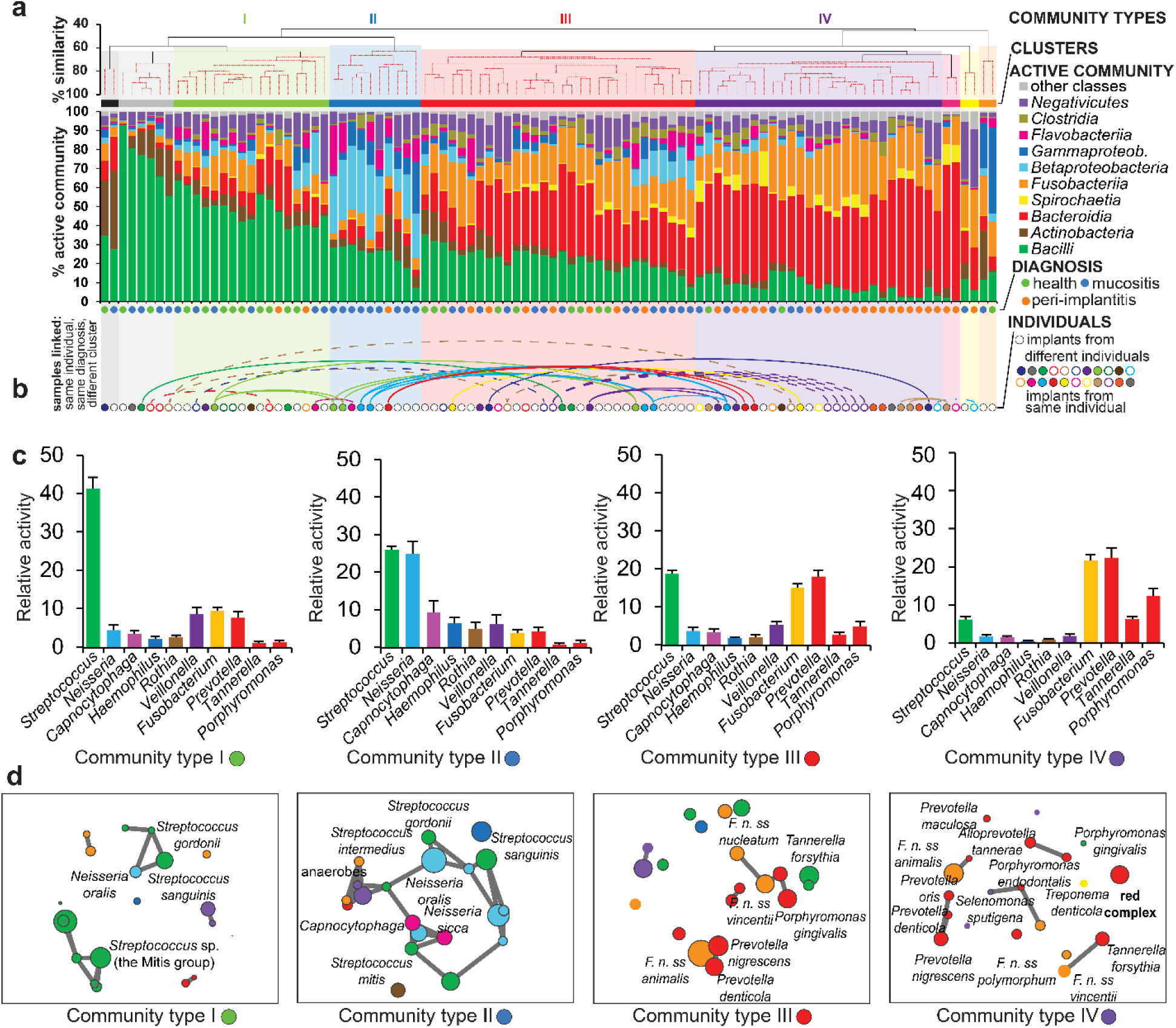
Major active community types at dental implants. **(A)** Clusters of class-level activity identified by SIMPROF. The main clusters representing major active community types (CT) were numbered I to IV and color coded. Composition of active community was plotted for every sample. Diagnosis is given below. **(B)** Inter-patient diversity. Samples originating from the same individual highlighted and linked by lines if they are placed in different clusters in spite of the same diagnosis. **(C)** Most active genera. **(D)** Species correlation networks.

### Microbial biofilm changes in peri-implantitis trigger numerous catabolic and a few specific anabolic pathways

Next, we analyzed changes in transcriptome-level enzyme expression, categorizing them into Enzyme Commission (EC) numbers, which provide a numerical classification for enzyme-catalyzed reactions. Taxon-independent analyses offer the advantage of capturing microbial functions rather than focusing on specific taxa. Given the strong patient-specificity of peri-implant biofilms and the functional redundancy among microbes, this approach enables detection of consistent, cross-patient signals that may be missed by taxon-focused methods. RNAseq generated a total of 2.6 billion reads across 98 samples, with an average of 26 million reads per sample. The identified transcripts were linked to 1824 ECs, of which 36% had a relative abundance of at least 0.1% in at least one sample. The overall expression patterns in PI were distinct from both PIH and PIM (PERMANOVA *p* = 0.0001), whereas PIH and PIM comparison did not show significant differences in their EC based profiles (*p* = 0.488) (**Supplemental Fig. 4**). Using CAP and DEseq2 analyses, we identified ECs with a significantly elevated expression in individual diagnosis groups (**Supplemental Table 6**). We then mapped these diagnosis-group associated ECs onto KEGG pathways to assess their associations within specific KEGG modules or sequential reaction chains (**Fig. 4**). ECs positively associated with PIH or PIM, were – in comparison to PI – primarily linked to anabolic processes, including the synthesis of multiple amino acids, glycerolipids, pyrimidines, and purines, as well as to pathways involved in denitrification, the pentose phosphate cycle, glycolysis, and the citrate cycle. ECs positively associated with PI represented catabolic processes, such as degradation of multiple amino acids, acyloglycerols, and galactose. Additionally, some ECs involved in biosynthetic pathways, such as those for isoprenoids, purines, and serine, as well as carbon fixation and gluconeogenesis, were also positively associated with PI.

**Figure 4.**
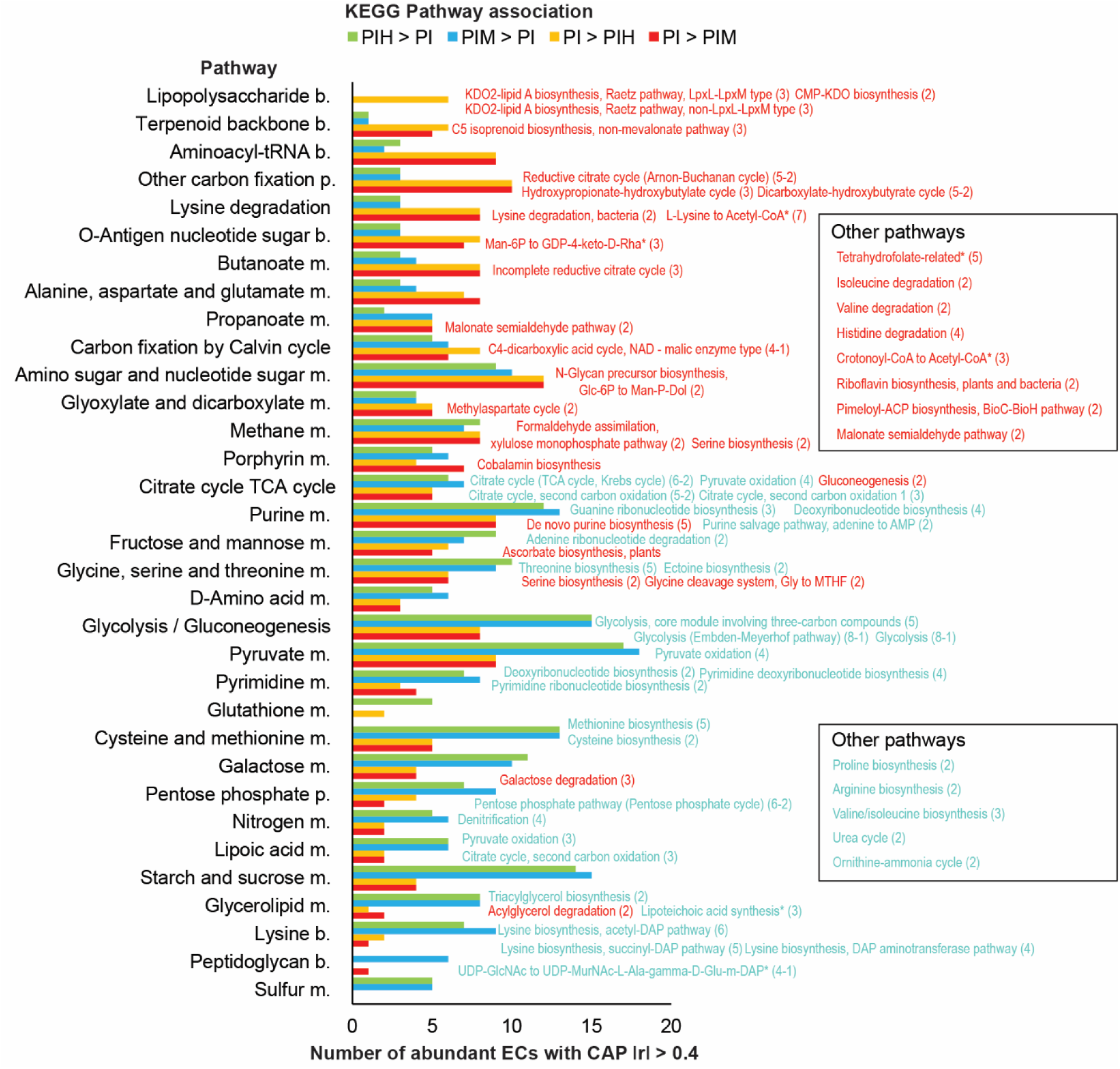
Pathways and modules associated with diagnosis groups. For each of the diagnosis groups, the bar plot displays the number of enzymes (ECs) from specific KEGG maps that were correlated with corresponding diagnosis using CAP. Individual KEGG modules are listed beside KEGG map bars if at least two members were associated with either PIH, PIM, or both (in cyan), or with PI in either or both comparisons (in red). PIH-PI and PIM-PI yielded largely overlapping results for both positively and negatively associated features with PI. In contrast, PIH-PIM showed no significant associations. Modules containing both PI-associated and non-PI-associated enzymes were excluded unless they were clearly dominated by one group (with a difference of at least three member ECs). In the brackets following each module name, the number of associated members is shown, if applicable minus the count from the opposite group. Consecutive reactions spanning three or more steps not assigned to a KEGG module but showing correlation are labeled as custom reactions, indicated by an asterisk.

PIM samples representing CT II and CT III differed by the expression of 219 ECs (*P* < 0.05, **Supplemental Table 7**). PIM CT II was mainly linked to pyruvate and lipoic acid metabolism, while CT III was characterized by lysine degradation and butanoate metabolism, which are functionally interconnected. Finally changes in amino sugar and nucleotide sugar metabolism was observed in both PIM-associated CTs. This data, along with biofilm biomass estimates derived from bacterial RNA yields, indicates that the difference between PIH and PIM is predominantly driven by biofilm expansion with minimal functional changes. In contrast, the PIM and PI differences involves both biofilm expansion and significant functional alterations.

### Key microbial players demonstrate class specific functions

To understand the physiology and functional roles of microbial members within the biofilm, we examined class-specific activities in peri-implant biofilms (**Fig. 5**, **Supplemental Table 8**). Ordination (**Fig. 5a**) and clustering (**Fig. 5b**) revealed unique activity fingerprints for each class, consistent across diagnosis types (**Fig. 5b**). Pairwise comparisons of the four most active bacterial classes (*P* < 0.05), identified class-associated ECs that were subsequently grouped into KEGG maps (**Fig. 5c**). Bacilli, associated with PIH/PIM (often representing CT I), exhibited higher activity in carbohydrate, nucleobase, and pyruvate metabolism, as well as the pentose phosphate pathway. Negativicutes, also PIH/PIM-associated, showed enriched nucleobase, porphyrin, lipopolysaccharide, and amino acid metabolism. Bacteroidia, PI-associated as well as abundant and highly active in CT III/CT IV, were characterized by amino sugar, purine, and one-carbon metabolism. Fusobacteria exhibited porphyrin metabolism, similar to Negativicutes.

**Figure 5.**
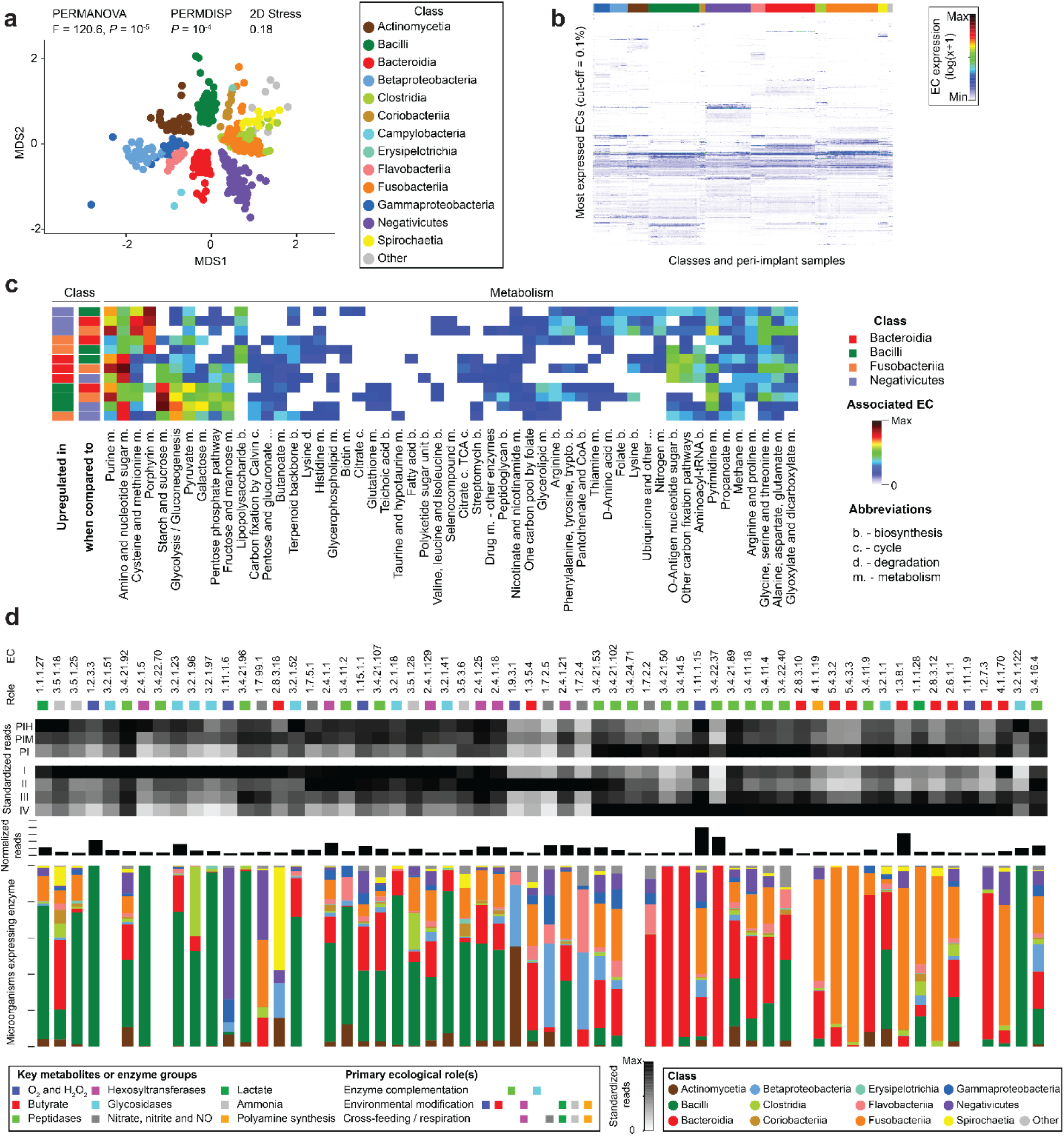
Unique functions of microbial classes in peri-implant biofilms. **(A)** Unconstrained ordination of EC-level expression data, with the microbial classes in each sample treated as a separate sample, resulting in n=541 pseudo-samples. MDS plot calculated on Bray-Curtis dissimilarities for log(x+1)-transformed read counts grouped to enzymes (E.C., relative activity cut-off = 0.1%, number of ECs = 1370). Datapoints are colored by class. **(B)** Shade graph for class activity across samples. Class-samplecombinations and enzymes were each sorted using hierarchical clustering on Euclidean distances. Same input data as for “A”. **(C)** Shade graph for class-specific activities. ECs which activity was associated with a given class were grouped to KEGG maps. **(D)** The top 60 most highly expressed ECs with predicted ecological relevance were analyzed. ECs were grouped based on Spearman correlations, and their expression levels were standardized by their maximum values and visualized as a heatmap across diagnostic groups and community types. Below, normalized counts and the composition of producers are illustrated. ECs are color-coded by ecological groups, defined by their associated metabolites, enzymes, and ecological roles. Producer data is presented at the class level.

To uncover the ecological functions of peri-implant microbial classes, we linked the ECs to their ecological roles and associated metabolites, analyzing the top 60 most highly expressed features across diagnosis groups and CTs (**Fig. 5d**, **Supplemental Table 9**). Glycosidases from Bacilli and peptidases from Bacteroidia, known to participate in enzyme sharing for synergistic degration of host derived polymers like salivary glycoproteins^31, 32^, showed the highest expression in PIH/PIM and PI, respectively, with β-galactosidase (EC 3.2.1.23) and Arg-gingipain (EC 3.4.22.37) being the most highly expressed enzymes. H_2_O_2_ production (EC 1.2.3.3), primarily driven by Bacilli and Negativicutes-driven H_2_O_2_ detoxification (EC 1.11.1.6) was observed in PIH and PIM. Among oxidative stress related enzymes, peroxidases (EC 1.11.1.15) showed the overall highest expression. Hexosyltransferases activities from Bacilli were indicative of carbohydrate polymer biosynthesis, while butyrate and polyamine production driven by Fusobacteria was associated with PI. Notably, the PIM CT II subtype exhibited unique activities, including oxygen depletion (EC 1.9.3.1), succinate/fumarate metabolism (EC 1.3.5.4), denitrification (EC 1.7.2.4/5), and glycogen synthesis (EC 2.4.1.21), highlighting distinct ecological adaptations in peri-implant biofilms.

### Bacteriophage activity clusters with different types of peri-implant disease

Despite their critical role in shaping bacterial populations and influencing biofilm physiology, the peri-implant phageome remains poorly characterized^33^. To replicate, bacteriophages must hijack the host bacterial machinery. As a result, MTX captures transcripts derived from active phages and phage-like elements, providing a functional readout of phageome activity. Using the IMG/VR database as a reference, we assessed transcription levels of phage-like genes, grouped as provisional species-like vOTUs (*i.e*., viral operational taxonomic units representing sequence bins). Ordinations based on phage data showed distinct separation between PIH, PIM, and PI samples (PERMANOVA *P* = 6 · 10^-5^). CAP and DEseq2 analysis revealed phage species (vOTUs) associated with each diagnosis group (**Supplemental Fig. 5**, **Supplemental Table 10**). Analysis of phage hosts revealed high activity of phages targeting typical oral microorganisms with distinct patterns across diagnosis groups. However, a significant fraction of phage activity remained unassigned to specific bacterial hosts (**Fig. 6a**). Phages targeting *Fusobacterium* and *Prevotella* exhibited higher activity in PI than in PIH or PIM, where phages targeting *Streptococcus* and *Haemophilus* were more active (**Fig. 6b**). Additionally, *Veillonella* phages showed greater activity in PIM compared to PI. The *Streptococcus* phage, vOTU 20193, exhibited the overall highest average relative activity of 6.6% across all samples (**Fig. 6c**). In conclusion, variations in the active phageome across diagnosis groups reflected changes in both the abundance and activity levels of phage hosts.

**Figure 6.**
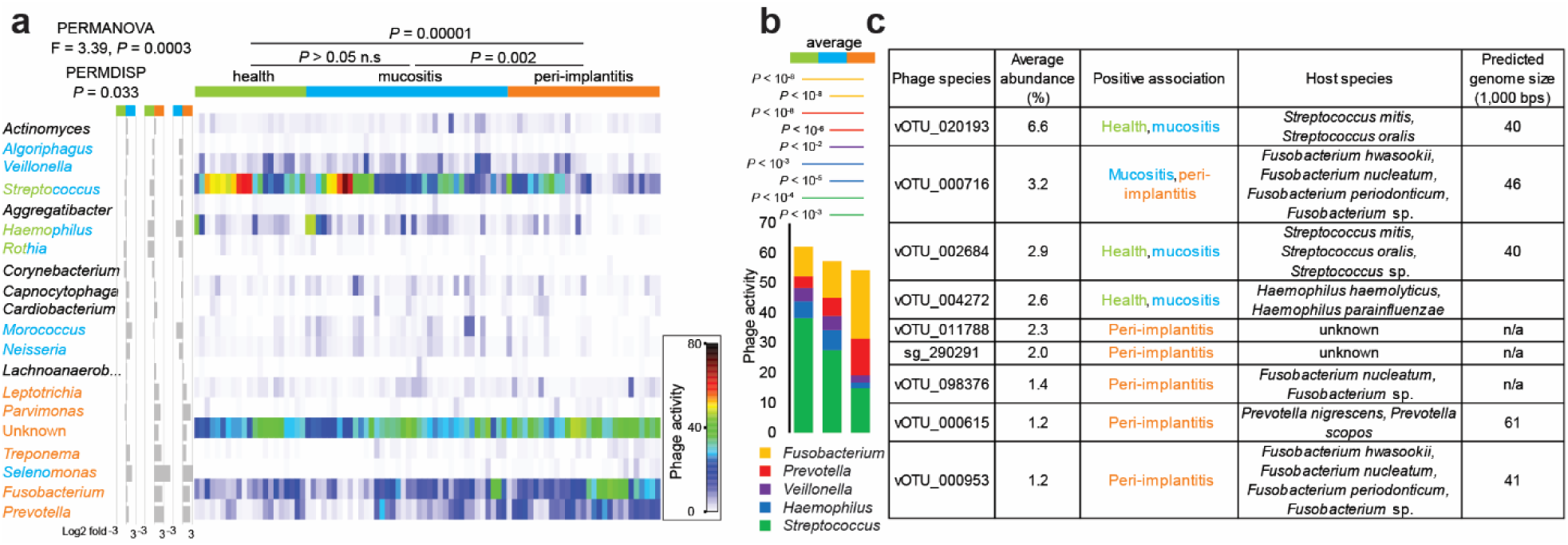
Active phageome across diagnosis groups. **(A)** Heatmap for phages grouped by a host. Diagnosis associations are indicated by font color. **(B)** The activity of phages infecting five genera dominated the biofilms and exhibited high diagnosis specificity. **(C)** Most abundant peri-implant phage species (average relative activity cut-off = 1%). Green, blue and orange colors represent peri-implant samples from health, mucositis and peri-implantitis, respectively.

### Peri-implant disease exhibits distinct host response to biofilm

Detection of intact human ribosomal RNA, indicating higher transcript integrity, enabled minimally invasive transcript-level analysis of host activity at the local host–microbiome interface We examined the expression of approximately 35,000 human genes (excluding low-count genes) around dental implants across diagnosis groups using DESeq2 and CAP analysis (**Supplemental Fig. 6**). The host gene expression patterns in PI differed significantly from both PIH (PERMANOVA *P* = 0.011) and PIM (PERMANOVA *P* = 0.004). In contrast, the comparison between PIH and PIM showed no significant differences in their profiles (*P* = 0.33). DESeq2 analysis revealed 102 differentially regulated genes when comparing PI and PIM with 32 genes upregulated and 70 downregulated in PI (DESeq2 adj *P* < 0.05) and the smallest number of differentially regulated genes in PIM vs. PIH comparison (3 genes upregulated in PIM). Using CAP analysis, we identified the top actively expressed genes (filtered at a 0.1% maximum cut-off) which showed a high correlation (ǀ r ǀ > 0. 25) with diagnosis groups. We further identified enriched GSEA hallmark gene sets, which represent specific well-defined biological processes and canonical pathways, associated with diagnosis groups (adjusted *P* < 0.05, tested for CAP outputs, **Fig. 7**). This analysis showed that keratinization was linked to PIH, while increased expression of genes relating to translational activity was characteristic for PIM compared to PIH. PIM samples representing CT II and CT III differed by the expression of only 12 host genes. Comparisons between PIH vs. PI and PIM vs. PI revealed similar host pathways associated with both PIH and PIM, primarily linked to keratinization and translation. An increased abundance of transcripts associated with hemidesmosome assembly was notably characteristic of PIM in the PIM vs. PI comparison. PI signature pathways were largely consistent across comparisons, predominantly associated with TNF-alpha signaling via NF-κB, inflammatory responses, cytokine signaling, interferon signaling and hypoxia. In summary, the host response in PIH and PIM was similar, reflecting an intact epithelial barrier and housekeeping functions, while PI was strongly associated with inflammation and related signaling.

**Figure 7.**
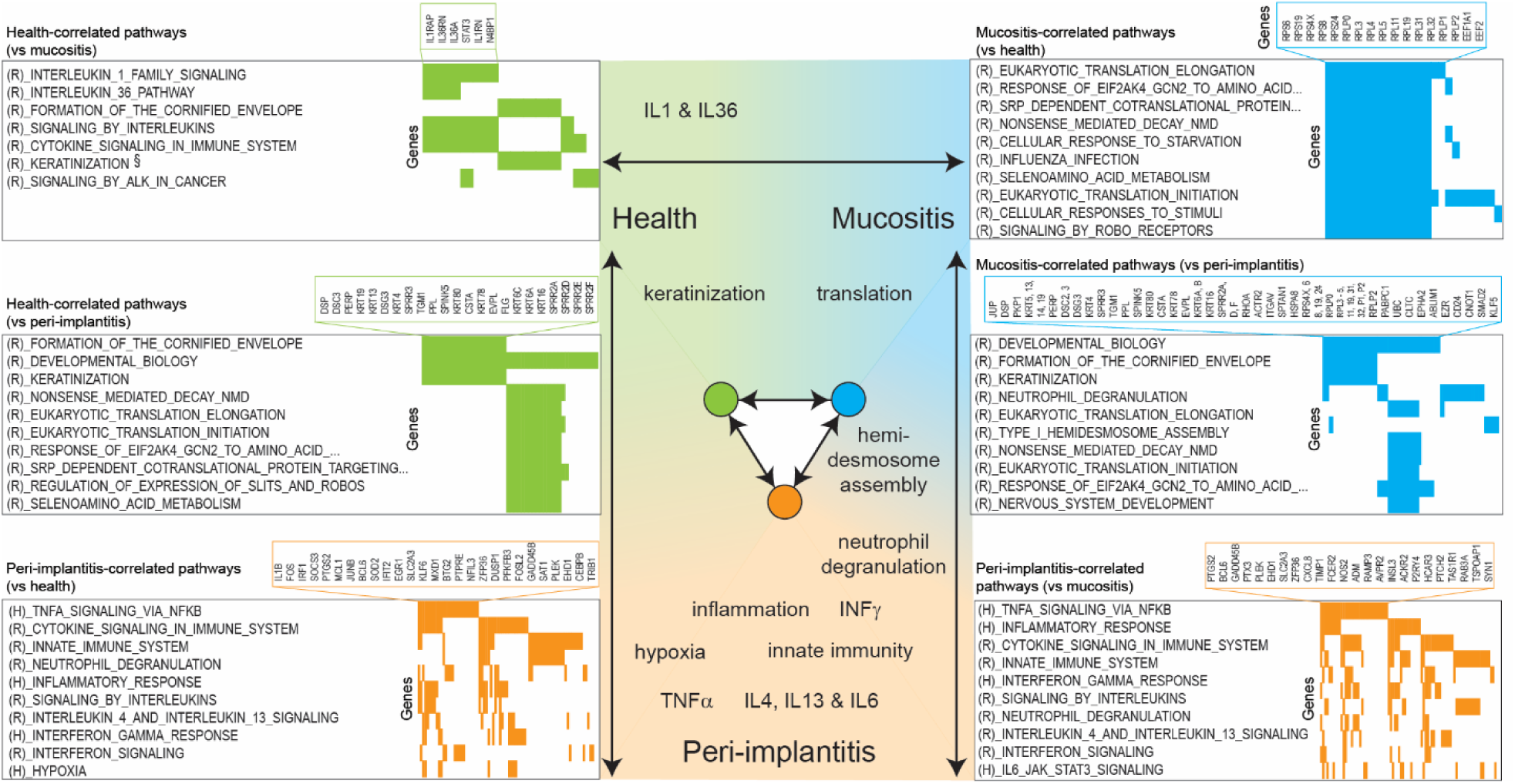
Host gene expression across diagnosis groups. Pathways representing genes correlated with diagnosis groups (ǀ r ǀ > 0.25) in CAP. All or top 10 pathways are listed for each comparisons and diagnosis group based on increasing adjusted *P* value q-value. Genes representing the top pathway are listed. (Ranges of adjusted p-values from top pathway (from top left graph, clockwise): 5.5 · 10^-5^ – 1.1 · 10^-2^, 3.2 · 10^-29^ – 1.0 · 10^-21^, 1.7 · 10^-30^ – 1.0 · 10^-14^, 3.2 · 10^-40^ – 5.1 · 10^-10^, 1.2 · 10^-32^ – 6.0 · 10^-10^, 2.0 · 10^-25^ – 1.5 · 10^-14^. (H) – Hallmark gene sets, (R) – Reactome subset of canonical pathways. Green, blue and orange colors represent peri-implant samples from health, mucositis and peri-implantitis, respectively.

### Multi-omic integrative analysis revealed complex interactions in the peri-implant ecosystem

We leveraged our multi-omics, multi-layered dataset to trace key interactions within the peri-implant ecosystem. by investigating pairwise interactions among bacterial species abundances (full-16S-based), enzymes, phages, and host genes (all MTX-based), using hierarchical association testing (**Fig. 8**. This was followed by integrative analyses combining features from all datasets.

**Figure 8.**
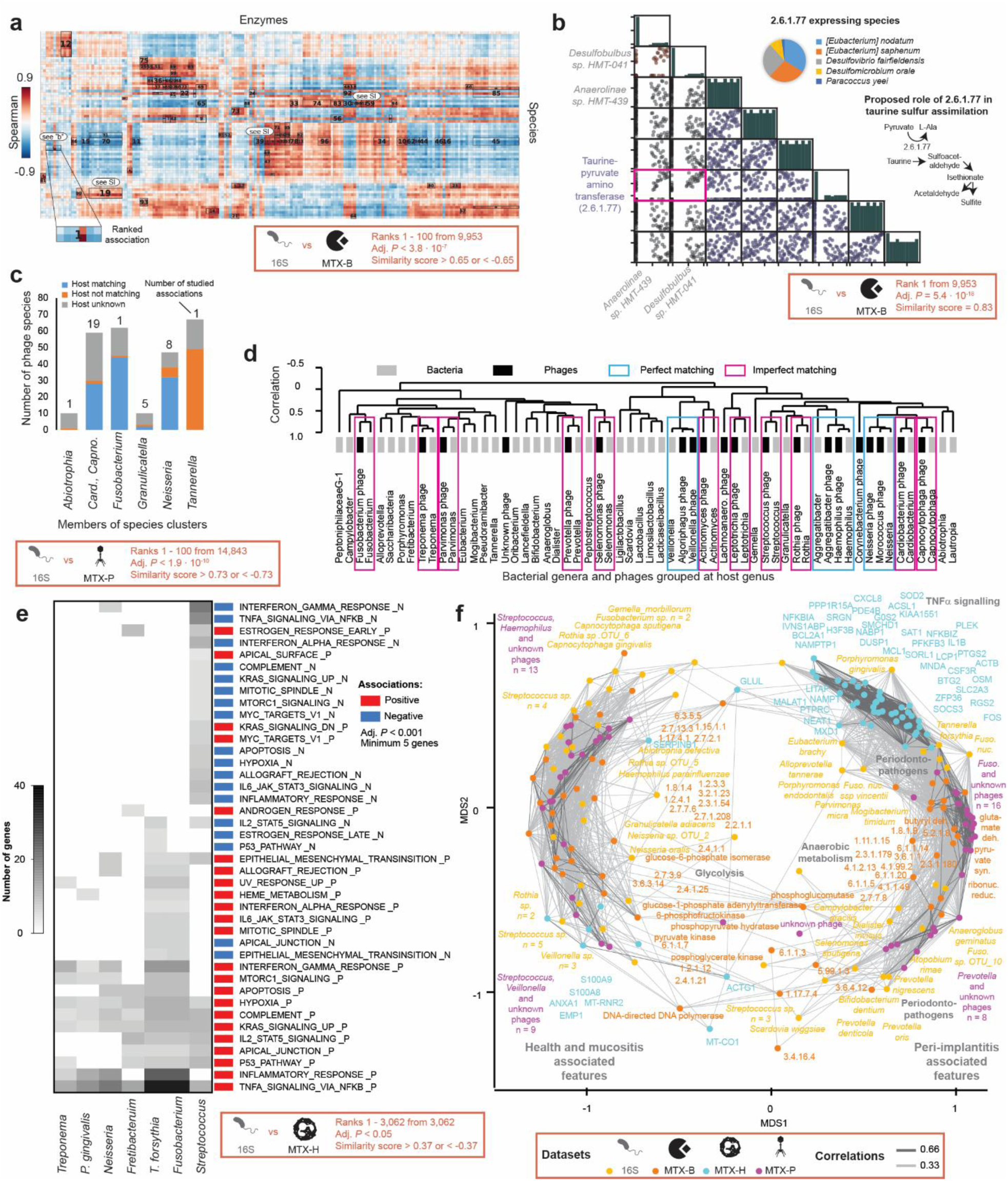
Network of interactions between microbial, ecological, biochemical, and host factors. **(A)** Associations between species clusters (at DNA level) and enzymes (at RNA level), with the top 100 associations displayed on the heatmap. **(B)** the strongest association from panel “A”. **(C)** relationship between phage-associated species and predicted hosts of those phages. **(D)** correlations between genera and phages grouped by their host genus. **(E)** Relationship between species (at DNA level) and host reponse (at RNA level). **(F)** Integrative clustering analysis of four datasets.

Profiling of species-enzyme relationships revealed potential interspecies metabolic dependencies (**Fig. 8a**, **Supplemental Table 11**). The strongest association, identified from approximately 10,000 statistically significant correlations, was a positive correlation between Anaerolineae sp. HMT-439, *Desulfobulbus* sp. HMT-041 and taurine–pyruvate aminotransferase (EC 2.6.1.77), an enzyme involved in taurine sulfur assimilation, predominantly expressed by *Eubacterium*, *Desulfovibrio* and *Desulfomicrobium* species (**Fig. 8b**). Thus, our integrative analysis identified distinct clusters of highly correlated species and enzymes, further distinguishing between active species that directly express these enzymes as shown from the EC transcripts assigned to them and other species that, despite lacking expression capabilities, are likely to be associated with or dependent on these activities.

The corresponding microbial taxa (full-16S) and bacteriophages (MTX) analysis identified 14,843 associations (**Supplemental Table 12**), from which top ranked associations were compared to identify potential phage hosts (**Fig. 8c**). Phages were typically linked to their host bacteria, except for *Fusobacterium* phages which were associated with *Tannerella forsythia*, suggesting it may benefit from phage-induced lysis. Interestingly, correlation between phage RNA reads aggregated by their hosts and bacterial genera at DNA level showed a near-perfect matching (**Fig. 8d**).

The microbial taxa (full-16S) and host response genes (MTX) analysis identified 7,482 associations (**Supplemental Table 13**). The top 10 associations comprised five clusters representing nine predominantly disease-associated microbial species and seven clusters encompassing 40 host genes (FDR-adjusted q < 7.06 10⁻⁷, Spearman’s ρ > 0.67). Unconstrained analysis was followed by an analysis of selected associations of representative key biofilm members (**Fig 8e**). 107 associations were found between representative taxa and host genes. They were grouped to 24 hallmark sets. *Streptococcus* exhibited a distinct profile characterized by a high number of negative associations. *Neisseria* grouped with disease-associated taxa, while *Fusobacterium* clustered closely with *T. forsythia*. Additionally, *P. gingivalis* displayed a similar profile to that of *Treponema*.

At functional level, we identified 36,617 associations between bacterial enzymes (MTX) and host response genes (MTX) (**Supplemental Table 14**). Gingipain R, regarded as one of the most important enzymes causally linked to periodontal and peri-implant pathologies^34^, in cluster with mannase (3.2.1.25), was positively associated with 61 host gene clusters representing TNFα signaling (FDR-adjusted q = 6.5 · 10^-35^), inflammatory response and 12 other hallmark gene sets.

Integrative clustering analysis of the most abundant and active elements across four datasets consistently depicted the peri-implant ecosystem (**Figure 8f**). The major distinctions were observed between PIH/PIM and PI, which was evident at all molecular and taxonomic levels.

## Discussion

In this study, we utilized the largest clinical cohort to date with paired microbiome composition and metatranscriptomic profiling of implant-associated biofilms to integrate multiple molecular features and present the first high-resolution *in situ* characterization of the peri-implant ecosystem across peri-implant health (PIH), peri-implant mucositis (PIM), and peri-implantitis (PI). While overall population dynamics aligned with patterns observed in previous studies on microbiome composition ^7–9, 12, 14, 35–38^, our integrative approach, incorporating metatranscriptomics, revealed novel disease-associated features. These findings include absolute levels of biofilm activity, active community types, the activity of ecologically relevant enzymes and phages, as well as characteristic host response patterns. Furthermore, association testing provided a broader context by uncovering potential bacteria-enzyme, phage-bacteria, and microbe-host interactions across different molecular levels. These insights provide a deeper understanding of the physiology, ecology and virulence of peri-implant biofilms.

Both clinical and molecular measurements, including absolute biofilm activity, consistently showed that biofilm expansion is a primary driver of the PIM development around implants and contributes to its further progression to PI. Our findings revealed highly conserved enzyme expression among biofilm members, pointing to substantial functional specialization and interdependence. Combined with the high species diversity, these characteristics may promote strong growth synergies within the biofilm^39^. Additionally, observed variation in host response, including non-canonical response profiles within diagnostic groups suggest an important role in biofilm regulation^40, 41^.

At RNA level, PIM showed the presence of an alternative community type that emerged in CT III alongside the typical composition of known periodontopathogens in CT II. This community, here described for the first time, was characterized by a high activity of *Neisseria* spp., which are typically being considered low-virulent^42^. Unexpectedly, our correlation analysis suggests that these species elicited host responses comparable to those triggered by established periodontopathogens. While some DNA-based clinical studies associate *Neisseria* with PIH^11, 14, 36, 37^, others link it to PIM^6^ or less-severe PI^38^, or report no association but generally high abundance^43^, highlighting a potential dual role in oral health and disease. Several factors may regulate *Neisseria* populations. Bleeding, a hallmark of PIM, provides heme, potentially supporting *Neisseria* dominance^44^. However, under anaerobic conditions, *Neisseria*’s activity diminishes unless alternative electron acceptors, such as nitrate, are available, potentially sourced from diet or biofilm metabolism^45, 46^. This dynamic could explain the presence of CTs with or without high *Neisseria* activity. Our findings suggest that *Neisseria* may contribute to the early peri-implant pathology in patient subpopulations. Further investigation into host-*Neisseria* interactions in the peri-implant environment is warranted to better understand their potential role as a pathobiont. Additionally, experimental^47^ or observational^48^ longitudinal clinical studies could provide further insights into whether *Neisseria* activity modulates the progression dynamics of mucositis.

For the first time, we specifically analyzed the expression of enzymes previously reported or predicted to be ecologically relevant^31^ aiming to assess their functional significance for the peri-implant ecosystem. As expected, key enzymes were involved in oxidative stress generation or reduction, enzyme sharing, and metabolic cooperation. While we confirmed several important contributors^49–53^, we also identified ecologically active enzymes and taxa that have not yet been included in existing interaction models and warrant further investigation. Community type- or disease state-specific patterns revealed potential target interactions for anti-biofilm interventions. Beyond bacterial activity, the peri-implant phageome emerges as a significant yet understudied^33^ component of peri-implant biofilms. In the present study, we uncovered a remarkable diversity of active oral phages and phage-like elements within the peri-implant ecosystem, consistently identifying them as a key biofilm constituents across diagnostic groups and microbial communities. The strong association between *Tannerella forsythia* and *Fusobacterium* phages suggests that phage-induced lysis may facilitate the expansion of neighboring fastidious species. These findings are supported by previous experiments showing that conditioned media from *Fusobacterium* spp. cultures promote the growth of fastidious microorganisms, including *Prevotella* spp., phylogenetically related with *Tannerella* spp.^54^. Additionally, the activity of phages with unknown hosts correlated to the presence of specific bacterial species, thus uncovering potential hosts such as *Capnocytophaga* and *Fusobacterium* spp. Our approach could be applied to refine current host assignment methods for phages in the future.

Although plaque-derived host RNA may originate from exfoliated or lysed epithelial and immune cells, it reflects the host-microbiome interface in situ and provides valuable, minimally invasive insight into localized immune responses. This approach has been increasingly recognized in respiratory microbiome studies^23, 24^, and exploratory analyses of host transcripts from oral samples such as saliva, buccal swabs, and gingival crevicular fluid have shown promise^55^. The comprehensive profiling of the host response across disease states associated distinct host activity patterns with each diagnostic group. Notably, upregulation of keratinization in PIH (vs. PIM) and in PIM (vs. PI), emphasizes the critical role of keratinized epithelium in maintaining peri-implant health. Keratinized epithelium protects mucosal tissues from microbial insults, reducing vulnerability and tissue damage^56^. Insufficient keratinized mucosa around dental implants has been linked to increased plaque accumulation, tissue inflammation, mucosal recession, and attachment loss^3, 57^. These findings suggest that the stability of soft tissues surrounding osseointegrated dental implants significantly impacts bone regenerative processes and long-term outcomes. PI was strongly associated with inflammatory responses, such as TNF-α signaling via NF-κB, cytokine signaling, interferon signaling, and hypoxia. TNF-α is a key mediator of periodontal inflammation and alveolar bone resorption^58, 59^. The detection of host gene expression related to hypoxia in PI corresponds with the observed dominance of anaerobes in biofilms, which are major producers of lipopolysaccharides (LPS)^60^. The hypoxic environment may enhance TNF-α and other cytokine expressions in response to biofilm-derived LPS, subsequently activating the NF-κB pathway^61^. Together, these molecular signatures offer a coherent and detailed understanding of the pathological mechanisms underlying PI.

Our integrative analysis revealed an intriguing link between the PI-associated *Anaerolineae*-*Desulfobulbus* cluster and taurine–pyruvate aminotransferase, which is involved in anaerobic respiration in *Eubacterium*, *Desulfovibrio*, and *Desulfomicrobium*, as the strongest association. Taurine, a ubiquitous sulfonate metabolite, accumulates in inflammatory lesions^62, 63^. *Anaerolineae* and sulfate-reducing bacterium ‘SRB’ *Desulfobulbus* have been associated with periodontal disease^64–69^, produce virulence factors^54, 70^ and rely on scavenging essential metabolites, due to the loss of some biosynthetic capabilities^70–72^. This suggests that taurine-metabolizing *Eubacteriu*m spp. and two SRBs (*Desulfovibrio* and *Desulfomicrobium*) may provide growth factors or aid in the removal of inhibitory metabolites for the *Anaerolineae*-*Desulfobulbus* consortium. These fastidious and sensitive SRBs have also been linked to disease^73, 74^ ^75, 76^ and may depend on *Anaerolineae*, *e.g.*, for the breakdown of complex organic compounds or the production of growth factors, as a similar interactions were observed in gut for SRBs^77^.

Although our study is the first to provide integrative insights into key molecular interactions in peri-implant health and disease and drivers of dysbiosis by using multi-omics, also certain limitations should be considered, such as the lack of spatial information to measure ecological interactions ^78^. While our sample size is the largest in peri-implant metatranscriptomics to date, longitudinal studies are needed in the future to improve the understanding of biofilm dynamics, validate causal effects and identify population-specific responses,. Nevertheless, our integrated multi-omics approach provides a robust framework for future ecological and clinical studies, with the ultimate goal of improving diagnostic, preventive and treatment strategies for peri-implant disease.

## Materials and Methods

### Study design and ethical statement

The present clinical cross-sectional study was conducted at the Department of Prosthetic Dentistry and Biomedical Materials Science, Hannover Medical School, Germany, as part of the ‘SIIRI Biofilm Implant Cohort (BIC)’. SIIRI (Safety Integrated and Infection Reactive Implants) is an interdisciplinary initiative that aims to recruit hundreds of individuals and monitor their peri-implant microbiome over a period of up to 12 years to investigate the development of peri-implant diseases and devise early detection and prevention strategies. The study protocol was approved by the institutional ethics committee of the Hannover Medical School, Germany (No. 9477). Written informed consent was obtained from all participants that had at least one dental implant diagnosed with either peri-implant health (PIH), peri-implant mucositis (PIM) or peri-implantitis (PI) based on the 2017 World Workshop on the classification of periodontal and peri-implant diseases and conditions^29^. Detailed disease definitions and clinical examination as well as inclusion and exclusion criteria are described in **Supplementary material**.

### Sample collection and co-isolation of DNA and RNA

Peri-implant biofilm samples were collected from six sites per implant using sterile paper points (ISO 35/2.0, VDW GmbH) and curettes (HuFriedy Mfg. Co. LLC) to maximize input mass^79^. Samples were pooled in RNAprotect (Qiagen), incubated for at least 5 minutes, and stored at -80°C until further processing. DNA and RNA were co-isolated using a published protocol ^80, 81^ with modifications. The sample collection and co-isolation procedures are described in **Supplementary material**.

### Full 16S rRNA gene amplicons sequencing and RNA sequencing

Full-length 16S rRNA gene amplicon sequencing (full-16S) was performed on a PacBio Sequel system, followed by bioinformatic analysis using an in-house pipeline^82^. The dataset included 125 samples with a total of 2,012,347 reads, with an average sequencing depth of 13,783 reads per sample.

For metatranscriptomic analysis (MTX), ribosomal RNA was depleted using the Ribo-Zero Kit Epidemiology, and mRNA sequencing was performed on an Illumina HiSeq 2500 (50 or 68 base pairs). The quality filtered sequences were mapped to a custom oral metagenome reference created from eHOMD^30, 83^ and Pasolli et al.^84^ and further annotated with eggNOG^85^. To characterize functional features, counts were aggregated based on gene-EC relationships, and taxonomic information^86^. Phageome was characterized using IMG/VR database as reference^87^. Of 2.7 billion raw reads, 40% mapped to the human genome, and 1.4 billion were classified as bacterial or viral. 350 million mRNA reads had EC annotation assigned. The details of the full-16S and metatranscriptomic analyses are described in **Supplementary material**.

### Statistical analysis and association testing

Multivariate analyses were performed using PRIMER v7 with PERMANOVA+^88, 89^. Bray-Curtis dissimilarities or Euclidian distances of untransformed or transformed relative abundances were used for hierarchical clustering (group-average linkage, when applicable combined with SIMPROF permutation tests) and ordinations (nMDS, PCoA). Variables were compared across groups using PERMANOVA followed with testing for multivariate dispersion with PERMDISP. To account forinter- and intra-patient variability, PERMANOVA and PERMDISP were performed on profiles averaged across implants with the same diagnosis from the same patient. Diversity indices were calculated using DIVERSE. Canonical analysis of principal coordinates (CAP) was used to identify features correlated with disease groups. Differential expression analysis was conducted using DESeq2^90^ and controlling for patients, with significant features (log2 fold-change >1, adjusted p < 0.05). Functional pathway analysis was performed using KEGG Mapper for bacterial enzymes^91^ and Gene Set Enrichment Analysis (GSEA) for host genes^92^. For univariate comparisons, the Kruskal-Wallis test was applied with post-hoc Dunn’s or Wilcoxon Rank Sum test followed by Benjamini-Hochberg correction. For integrative analysis across 16S sequencing (species-level), metatranscriptomic bacterial (MTX-B), host (MTX-H), and phage (MTX-P) datasets, samples were standardized, filtered to retain features >0.1% in at least one sample, and log(x+1)-transformed. Significant associations (FDR-adjusted p < 0.05) were identified using hierarchical all-against-all association testing (HAllA)^93^.

## Data availability

The raw data and the supplementary data files supporting the findings of this study will be made available by the corresponding author upon request. All raw sequencing data are deposited to ENA under the Bioproject ID: PRJNA119296, which could be accessed after publication of the manuscript. STORMS checklist for this study could be accessed via https://doi.org/10.5281/zenodo.15495741

## Acknowledgments

This study was funded by the Deutsche Forschungsgemeinschaft (DFG, German Research Foundation) – SFB/TRR-298-SIIRI – Project-ID 426335750 (SPS, SH, and MSti). Additional support for this study was provided by the DFG under Germany’s Excellence Strategy (EXC 2155, Project Number 390874280, awarded to SH and MSti). WN, SH and MSti would like to thank Ministry of Science and Culture of Lower Saxony (Niedersächsisches Ministerium für Wissenschaft und Kultur) for funding through BacData, ZN3428. WB and IY are funded by the “Federal and State Program Promoting Female Professors”, Grant No. 01FP19068J. We also thank Marly Dalton and Rainer Schreeb for their valuable technical assistance. We thank Claudia Davenport for carefully reading and revising the manuscript.

