## supplementary material for "High-resolution taxonomic profiling and metatranscriptomics identify microbial, biochemical, host and ecological factors in peri-implant disease"

Short title: Peri-implant ecosystem; SI

Szymon P. Szafrański[1,2,3,$,*], Amruta A. Joshi[1,2,*], Matthias Steglich[1,2], Ines Yang[1,2], Taoran Qu[1,2], Wiebke Behrens[1,2], Uthayakumar Muthukumarasamy[4], Damianos Melidis[5], Paula Schaefer-Dreyer[1], Jasmin Grischke[1], Jan Hegermann[6], Wolfgang Nejdl[5,7], Susanne Häussler[3,4,8,9], and Meike Stiesch[1,2,3,$]

[1]Department of Prosthetic Dentistry and Biomedical Materials Science, Hannover Medical School, Hannover, Germany

[2]Lower Saxony Centre for Biomedical Engineering, Implant Research and Development (NIFE), Hannover, Germany

[3]Cluster of Excellence RESIST (EXC 2155), Hannover Medical School, Hannover, Germany

[4]Department of Molecular Bacteriology, Helmholtz Centre for Infection Research, Braunschweig, Germany

[5]L3S Research Center, Leibniz University Hannover, Hannover, Germany

[6]Research Core Unit Electron Microscopy, Institute of Functional and Applied Anatomy, Hannover Medical School, Hannover, Germany

[7]Knowledge-based Systems Laboratory, Leibniz University Hannover, Hannover, Germany

[8]Institute for Molecular Bacteriology, Twincore, Centre for Clinical and Experimental Infection Research, Hannover, Germany.

[9]Department of Clinical Microbiology, Copenhagen University Hospital - Rigshospitalet, Copenhagen, Denmark

*Szymon P. Szafrański and Amruta A. Joshi contributed equally

### Supplementary Methods

#### Clinical diagnosis and examination

Peri-implant health and disease diagnosis was based on the 2017 World Workshop on the Classification of Periodontal and Peri-implant Diseases and Conditions[1]. Accordingly, peri-implant health (PIH) was characterized by the absence of the following: clinical signs of inflammation, bleeding and/or suppuration, increased probing depth (PD) compared to previous examinations, or bone loss beyond crestal bone level changes due initial bone remodeling. Peri-implantitis mucositis (PIM) was diagnosed by the presence of bleeding and/or suppuration on gentle probing without the evidence of bone loss. The predefined criteria for peri-implantitis included presence of bleeding and/or suppuration on probing, increased PD and radiographic signs of bone loss as compared to previous examinations, or PD ≥6 mm and bone loss ≥3 mm in absence of previous examinations. The inclusion criteria for this study involved patients over 18 years of age who were systemically healthy, had at least one implant diagnosed with either PIH, PIM or PI. Patients with presence of periodontal disease, uncontrolled diabetes, heavy smoking (>20 cigarettes/day), recent use of antimicrobials or anti-inflammatory medications (within the last six months), pregnancy or lactation, conditions requiring pre-medication before dental procedures, or prior treatment for peri-implant disease on the same implant (except for supragingival prophylaxis) were excluded.

A single experienced clinician recorded detailed medical and dental histories and conducted chair-side examinations for all participants. The patient-level information including age, gender, history of periodontitis and smoking status were recorded, followed by full mouth examination to assess presence of periodontitis, residual teeth and oral hygiene maintenance. For each implant included in the study, clinical parameters including bleeding on probing (BOP), peri-implant probing depth (PD), suppuration, gingival index (GI), plaque index (PI), periotron values type of superstructure (fixed or removable) and duration of implant in function were recorded. Population-specific demographics and implant-specific clinical characteristics are presented in **Supplemental Table 1** and **Supplemental Fig. 1**.

**Biofilm sample collection and DNA-RNA co-isolation**

To efficiently collect biofilm samples and obtain both DNA and RNA from low-biomass samples while preventing RNA degradation and ensuring effective biofilm cell lysis, a tailored column-based protocol was employed, adapted from a previously published method[2, 3]. For sampling, sterile paper points (ISO 35/2.0, VDW GmbH, München, Germany) were inserted into the peri-implant sulcus or pocket at six different surfaces (mesio-buccal, mesial, mesio-lingual, disto-lingual, distal and disto-buccal) for 30 sec each, after isolating and drying the cervical region using cotton pellets. Additionally, subgingival plaque was collected at these sites using periodontal curette (HuFriedy Mfg. Co. LLC, Chicago, USA) and was transferred to the paper points before pooling them for each implant. Samples were incubated in RNAprotect (Qiagen, Hilden, Germany) for 5 min at room temperature. The samples were then immediately stored at -80 °C until further processing.

RNA isolation from the biofilm samples was performed as per previous protocol[2]. The protocol was modified to include co-isolation of DNA from the same samples for full-length 16S rRNA gene amplicon sequencing. The optimized protocol addressed key challenges associated with low-biomass biofilm samples, such as efficient lysis and prevention of RNA degradation as well as contamination. Chemicals were obtained from Sigma-Aldrich (Sigma-Aldrich, Taufkirchen, Germany) and the kits were used according to the manufacturer's instructions, if not stated otherwise. All glassware and other instruments were RNase-decontaminated using RNase ZAP solution (Ambion, Austin, TX, USA). The paper points were thawed and shredded with sterile scissors. The fragmented paper points were incubated in lysis buffer containing 10 mM Tris, 1 mM EDTA, pH 8.0, 2.5 mg/ml lysozyme and 50 U/ml mutanolysin at 25 °C for 1.5 h on a shaking incubator at 350 r.p.m. A total 700 µl of fresh buffer RLT (Qiagen) containing 1% (v/v) β-mercaptoethanol was added and vortexed for 10 sec. Samples (including the fragmented paper points) were placed on a QIAshredder Mini Spin column (Qiagen) and centrifuged for 1 min at 11,000 r.p.m. The flow-through containing the bacterial cells was mixed with 150 mg acid-washed and autoclaved glass beads (diameter 106 µm). Samples were vortexed 10 times for 30 sec at full speed with at least 1-min intervals on ice in between vortexing. The samples were then centrifuged for 1 min at maximal speed. Total RNA was isolated from the supernatants using the RNeasy Mini Kit (Qiagen). DNA was removed by column digestion and by DNase digestion in the eluate using the RNeasy cleanup procedure. Once the eluate for RNA extraction had been separated, DNA isolation was performed by placing the RNAeasy minispin columns in new tubes and incubated with 30 µl of 8 mM NaOH at 55 °C for 10 min. Eluate was collected by centrifugation and 3.03 µl 0.1M HEPES was added to it and both total RNA and DNA were stored at -80 °C until further use.

Blank controls were included to detect nucleic acid contamination due to reagents and paper points. Nucleic acid quality was assessed using the Agilent 2100 Bioanalyzer. To meet sequencing requirements, RNA samples with low yield but from the same patient and with same diagnosis were pooled, resulting in 24 pooled RNA samples from 55 healthy samples.

**Full 16S rRNA gene amplicons sequencing and analysis**

Full-length 16S rRNA gene amplicon sequencing (full-16S) was performed on a PacBio Sequel system, followed by bioinformatic analysis using an in-house pipeline[4]. In brief, the full-length 16S rRNA gene from each sample was amplified using the universal primer pair 27F (AGRGTTYGATYMTGGCTCAG) and 1492R (RGYTACCTTGTTACGACTT). PCR amplification was carried out with the KAPA PCR mix for 23 to 27 cycles under the following thermal conditions: an initial denaturation at 95 °C for 30 seconds, annealing at 55 °C for 30 seconds, and extension at 72 °C for 90 seconds, followed by a final elongation step of 10 minutes at 72 °C. After amplification, DNA concentration and quality were assessed using the Invitrogen Qubit dsDNA BR Assay Kit and the Qubit 2.0 fluorometer (Thermo Fisher Scientific). PCR products were then purified using the AMPure bead protocol, and samples with DNA concentrations of 5 ng or higher were selected for subsequent PacBio sequencing (PacBio Biosciences Inc., California, USA). SMRTbell libraries were constructed from the purified amplicons as per the manufacturer's protocol. Finally, circular consensus sequence (CCS) reads were generated from the raw PacBio data using the manufacturer's standard software suite.

PacBio CCS sequences were processed using an in-house pipeline[4, 5]. Taxonomic identification to the species level was achieved by comparing sequences against a modified bacteria-only version of the SILVA SSU Ref_NR 99 database version 132 [6], which was further enriched with Human Oral Microbiome

Database (HOMD)_16S_rRNA RefSeq Version 15.1 sequences[7], and the All-Species Living Tree Project (LTP) database version LTPs132_SSU[8], into which additional unnamed and phylotype sequences from HOMD 16S_rRNA RefSeq were additionally included. Taxonomic assignments were manually curated. Sequences that could not be confidently assigned to a single species were clustered into operational taxonomic units (OTUs) at 97% identity using UPARSE[9] as implemented in USEARCH 10.0.240. For higher-level taxonomic classification, the RDP classifier version 2.13 was employed at a cutoff bootstrap confidence of 80%[10]. To ensure data quality, sequences identified as potential contaminants, based on negative controls (blanks), literature information[11], and correlation analyses[12] were removed.

**RNA sequencing and analysis**

Eukaryotic cytoplasmic and mitochondrial ribosomal RNA, along with bacterial ribosomal RNA were removed using specific depletion probes and magnetic beads provided in the Ribo-Zero Kit Epidemiology (Illumina, San Diego, CA, USA). This step enhances the representation of messenger RNA (mRNA) in the sample by reducing the overwhelming abundance of ribosomal RNA, which typically constitutes over 90% of total RNA. The total RNA, enriched mRNA and cDNA were assessed using the 2100 Bioanalyzer system with RNA 6000 Pico and High Sensitivity DNA kits (Agilent Technologies Inc, Santa Clara, USA). Libraries were generated using the ScriptSeq v2 RNA-Seq (Illumina), which enables directional (strand-specific) library construction, preserving transcriptional orientation. Sequencing was performed in single-end mode (50 or 68 base pairs) on an Illumina HiSeq 2500 platform using the TruSeq SBS Kit v3—HS (Illumina).

BCL files were converted to FASTQ files (single-end reads) using the bcl2fastq conversion software version v2.20.0.422 (Illumina). Quality control was performed with fastp, a FASTQ preprocessor[13]. To distinguish human from non-human reads, we utilized a mapping approach incorporating bwa-samse (bwa-0.7.17, build 0.7.17-r1188[14]) aligning the reads to the human reference genome sequence GRCh38.p13 (GCA_000001405.28), as provided by the NCBI RefSeq, PRJNA31257, currently maintained by the Genome Reference Consortium. Instead of discarding the human-aligned reads, they were separated and retained for downstream analysis of host gene expression. Only unmapped reads were retained for downstream microbial or metatranscriptomic analyses, ensuring high specificity in non-human signal detection.

To construct a metagenomic reference set tailored to the human oral cavity, sequences from two publicly available datasets were selected and integrated into a single comprehensive database. The first source was the expanded Human Oral Microbiome Database (eHOMD, Rev: 2021-10-01, full eHOMD contig set), a well-established resource in the field of oral metagenome analysis, which includes 2087 bacterial genomes from the human mouth and aerodigestive tract[15]. The second dataset originated from the study of Pasolli *et al*. 2019[16], comprising 9,428 metagenomes and 154,723 metagenome-assembled genomes (MAGs) grouped into species-level genome bins (SGBs). From this source, the 7897 MAGs found in the study's oral-cavity-derived samples were selected. The sequences from both sources were combined into a combined oral metagenome reference set, grouped into 95% average nucleotide identity (ANI) taxonomic units (TUs) representing the species level, and annotated with the Prokka pipeline[17]. To extend the annotation and to assign additional EC numbers for further analysis, the eggnog annotated orthology database was applied[18]. The original taxonomy provided by eHOMD, along with Prokka and eggNOG annotations, together constituted the functional and taxonomic framework of the reference set. To ensure relevance to the study cohort, only TUs with sufficient coverage by sequencing reads from the collected oral metagenome samples were retained, determined through mapping using BWA-samse [15, 19]. For the final mapping of the sample reads, all rRNA-coding genes were excluded from the reference based on the associated annotations.

To quantify transcript abundance, sample reads were mapped to the protein-coding gene sequences from the selected TUs (SGBs) within the customized human oral reference set using BWA-samse. To avoid a biased counting for the transcript abundance of features, the reads mapped to multiple features were randomly distributed between them during the mapping of the sample reads against the customized human oral bacteria reference. Read counts were then extracted from the aligned sequencing read files using the htseq-count program from the HTseq package[20]. To streamline analysis and emphasize functional interpretation, the counts were aggregated based on their associated EC numbers. For genes annotated with multiple EC numbers, counts were summed at the highest hierarchical EC level where annotations were consistent. If a gene's EC annotations diverged at the second-highest hierarchical level, the corresponding read counts were categorized as "Unknown."

Taxonomic assignment of transcripts grouped by EC was performed following a method adapted from a previously published metatranscriptomic study[21]. Unlike the Jorth study, which reported species-level predictions, we opted to report genus- and class-level predictions to enhance accuracy. Briefly, read counts per locus were pre-filtered to include only the 1 million most abundant genes, in order to focus on biologically relevant transcripts and reduce noise from low-expression features. Hypothetical annotations were deselected and only genes with EC numbers were selected. For each sample, the counts per locus were normalized by the sum of read counts for all loci. Based on the functional annotation and taxonomic information of the individual gene, counts were then aggregated by the combination of EC number and taxon.

A total of 48,905 human-associated phage genomes from the oral cavity were selected from the IMG/VR database (IMG_VR_2018-07-01_4 – IMG/VR v2), based on metadata information in the IMGVR_all_Sequence_information.tsv file. These genomes were downloaded, indexed, and used as a reference for mapping RNA-seq reads using Bowtie2 (v2.3.5.1) with default settings. Read counts per genome per sample were used for the downstream multivariate analysis.

For the human transcriptome analysis, the latest annotated version of the human reference genome GRCh38.p13 (GCA_000001405.28), which includes 63,925 genes, was used as the reference. Samples with fewer than 500,000 reads mapped to human reference genome were excluded. To normalize for sequencing depth, read counts were standardized within each sample. Only the top 2,000 genes, accounting for ~70% of total reads on average, were retained for downstream analysis.

**Scanning Electron Microscopy**

Biofilm fragments were stored in fixative buffer containing 1.5% PFA, 1.5% GA, and 0.15M HEPES at 4°C. For dehydration, bacterial cells underwent a graded ethanol series treatment followed by desiccation with hexamethyldisilazane. The dehydrated samples were sputter-coated with gold, and imaging was performed using a Scanning Electron Microscope (SEM, Philips SEM 505) operating at 10 kV

### Supplemental Tables and Figures

**Supplemental Table 1 | Patient and implant characteristics.** Patient and implant characteristics were compared across three diagnosis groups: health (PIH) , mucositis (PIM) , and peri-implantitis (PI).

| | | | | | |
|---|---|---|---|---|---|
| **Demographic Characteristics** | | | | | |
| Total number of patients (n) | | 48 | | | |
| Total number of implants (n) | | 125 | | | |
| | | **PIH** | **PIM** | **PI** | ***p*-value** |
| **Population-specific parameters** | | | | | |
| Number of implants (n) | | 56 | 37 | 32 | *NA* |
| Patient age in years (Mean ± SD) | | 67 ± 10.04 | 67 ± 9.43 | 68 ± 8.15 | ns |
| Patient sex (n) | Females | 12 | 11 | 11 | ns |
| | Males | 10 | 13 | 6 | |
| Smoking (%) | Yes | 5.88 | 0 | 13.33 | ns |
| | No | 94.12 | 100 | 86.67 | |
| **Full mouth clinical parameters** | | | | | |
| Poor oral hygiene (%) | Yes | 14.28 | 14.28 | 46.6 | 0.02 |
| | No | 85.72 | 85.72 | 53.4 | |
| Residual teeth (%) | Yes | 95.45 | 100 | 88.23 | ns |
| | No | 4.54 | 0 | 11.77 | |
| **Implant -specific parameters** | | | | | |
| Gingival index (Mean ± SD) | | 0.22 ± 0.46 | 1.85 ± 0.57 | 2.35 ± 0.55 | < 0.0001 |
| Plaque index (Mean ± SD) | | 0.35 ± 0.61 | 0.92 ± 1.03 | 2.13 ± 0.84 | < 0.0001 |
| Implant position (%) | Incisor | 25.45 | 24.32 | 12.5 | ns |
| | Cuspid | 12.73 | 10.81 | 15.62 | |
| | Bicuspid | 34.55 | 21.62 | 34.38 | |
| | Molar | 27.27 | 43.24 | 37.5 | |
| Years of implant in function (Mean ± SD) | | 8.78 ± 6.96 | 8.96 ± 5.05 | 9.09 ± 4.96 | ns |
| Bleeding on probing (%) | Yes | 0 | 100 | 85.37 | < 0.0001 |
| | No | 100 | 0 | 14.63 | |
| Pocket depth (Mean ± SD) | | 2.79 ± 1.29 | 3.45 ± 1.32 | 7.13 ± 2.47 | < 0.0001 |
| Suppuration (%) | Yes | 0 | 0 | 53.13 | < 0.0001 |
| | No | 100 | 100 | 46.88 | |
| Periotron (Mean ± SD) | | 54.83 ± 39.36 | 56.79 ± 39.06 | 129.61 ± 45.54 | < 0.0001 |
| Pain (%) | Yes | 1.82 | 2.56 | 23.33 | 0.001 |
| | No | 98.18 | 97.44 | 76.67 | |

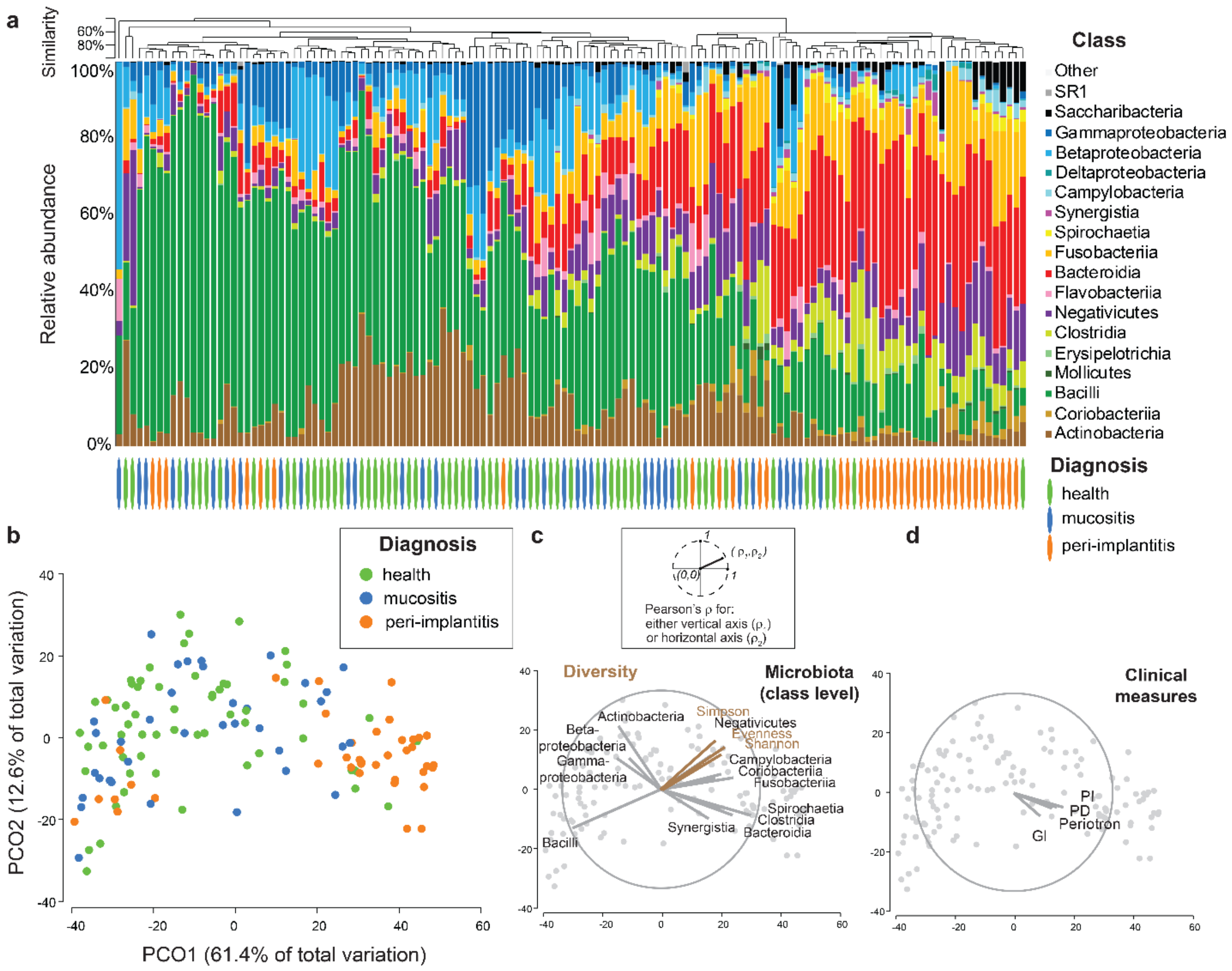


**Supplemental Figure 1. Composition of implant-associated biofilms at class level.** Taxonomy was assessed by full 16S rRNA gene amplicons sequencing. **a**, Composition of the 125 biofilms in 48 individuals. Biofilms were ordered by hierarchical clustering of Bray–Curtis similarity values calculated for non-transformed standardized class abundance profiles. Classes were grouped by phyla. **b**, Principal coordinates analysis (PCoA) of biofilm communities caluclated on data from „a”. **c**, Relationships between classes and diversity indices and ordination axes. D Relationships between clinical measures and ordination axes. Vector overlays in „c” and „d” were superimposed on the ordination from „b”.

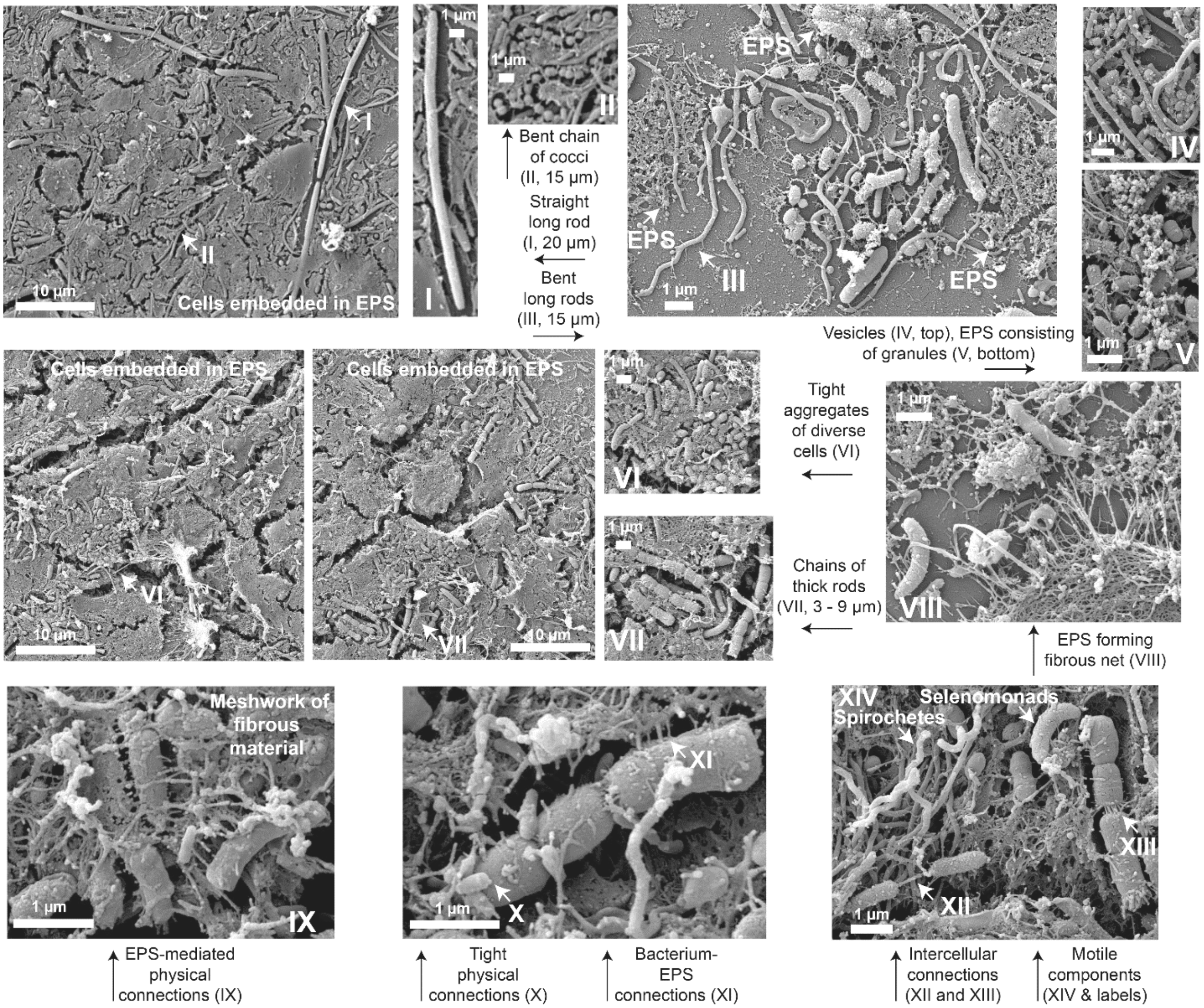


**Supplemental Figure 2. Biofilm morphology.** Scanning electron micrographs of submucosal implant-associated biofilms from peri-implantitis. Main components of biofilms are highlighted.

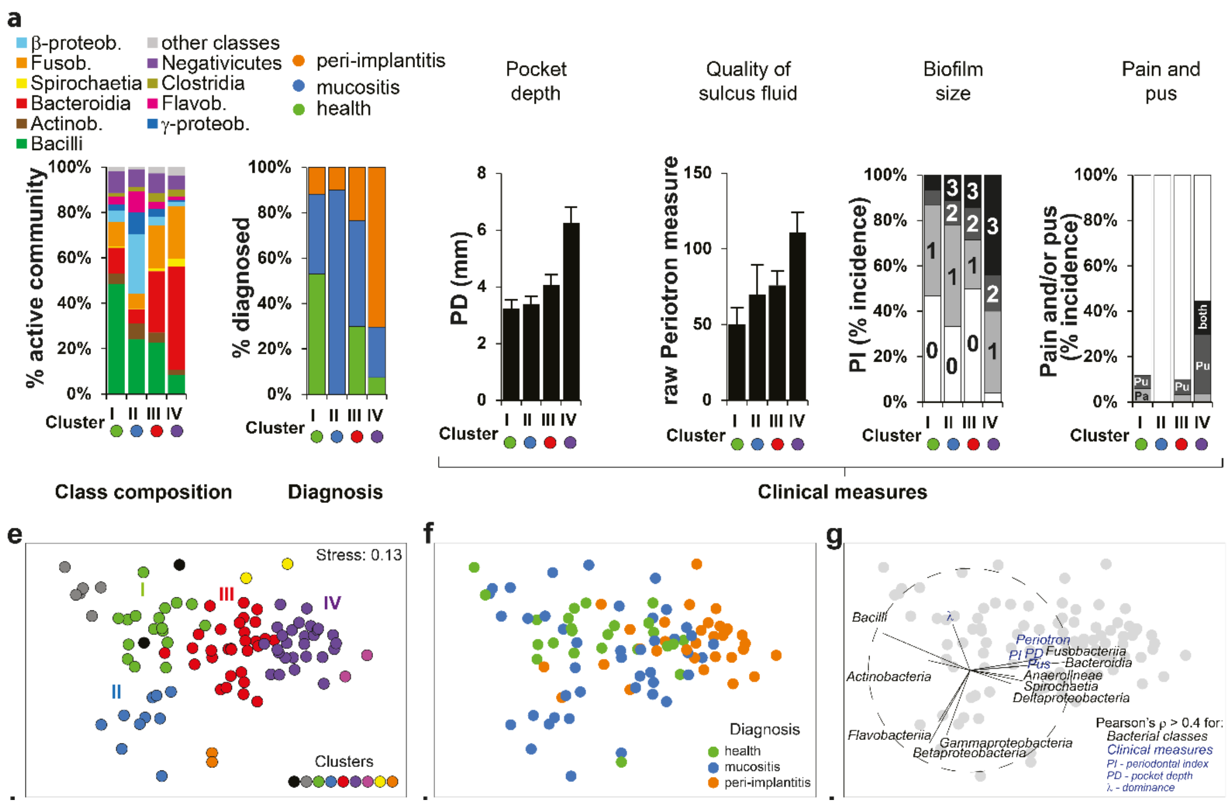


**Supplemental Figure 3. Four active community types. a**, General characteristics. Taxonomy at class level, diagnosis and four clinical measures are plotted for each community type. **b** – **d**, ordinations representing peri-implant samples overlaid with SIMPROF clusters, diagnosis, as well as classes and clinical measures as vectors, respectively. Non-metric multidimensional scaling (MDS) was used to represent the peri-implant samples in two-dimensional space.

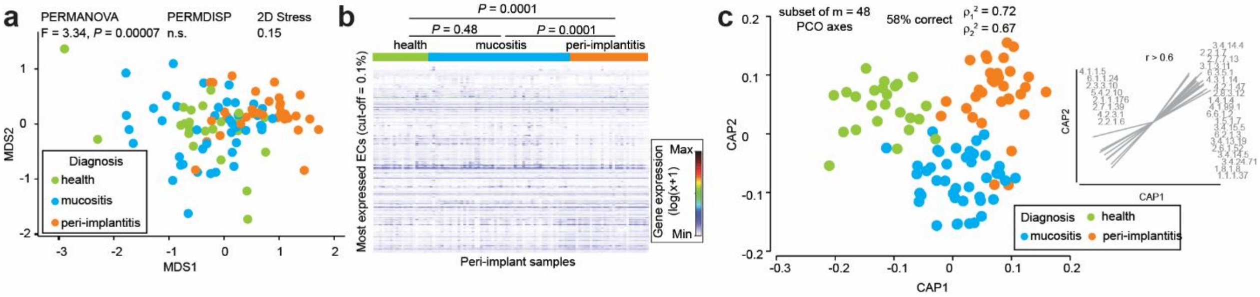


**Supplemental Figure 4. EC activities across diagnosis groups. a**, Unconstrained ordination for active ECs expression data. MDS plot calculated for X implants on Bray-Curtis dissimilarities for log(x+1)-transformed read counts grouped to ECs (relative activity cut-off = 0.1%). **b**, Heatmap for ECs. First, samples and ECs were each clustered, and next samples were sorted by diagnosis. **c**, Diagnosis-constrained ordination for ECs data. CAP plot calculated on same input data as for “a”. Selected diagnosis-correlated ECs ( | r | > 0.6) in CAP are indicated as vectors.

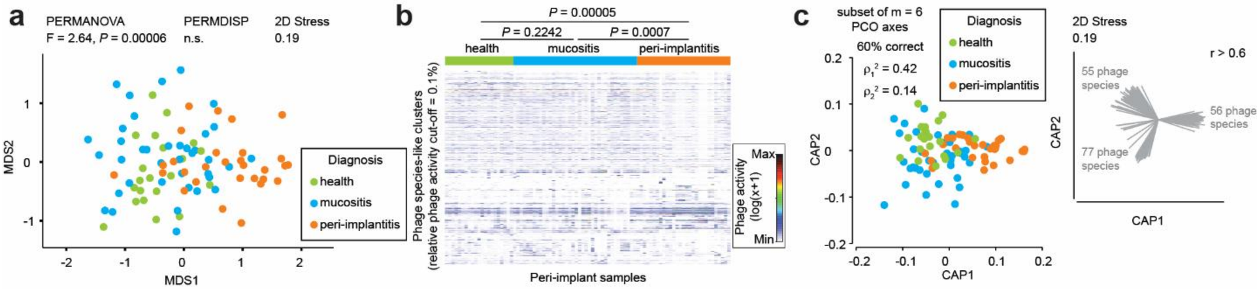


**Supplemental Figure 5. Active phageome across diagnosis groups.** **a**, Unconstrained ordination for active bacteriophage species (phage species, for short) expression data. MDS plot calculated for 88 implants on Bray-Curtis dissimilarities for log(x+1)-transformed read counts grouped to phage species (relative activity cut-off = 0.1%). **b**, Heatmap for phage species. First, samples and phage species were each clustered, and next samples were sorted by diagnosis. **c**, Diagnosis-constrained ordination for phage species data. CAP plot calculated on same input data as for "a". Selected diagnosis-correlated phage species ( | r | > 0.6) in CAP are indicated as vectors.

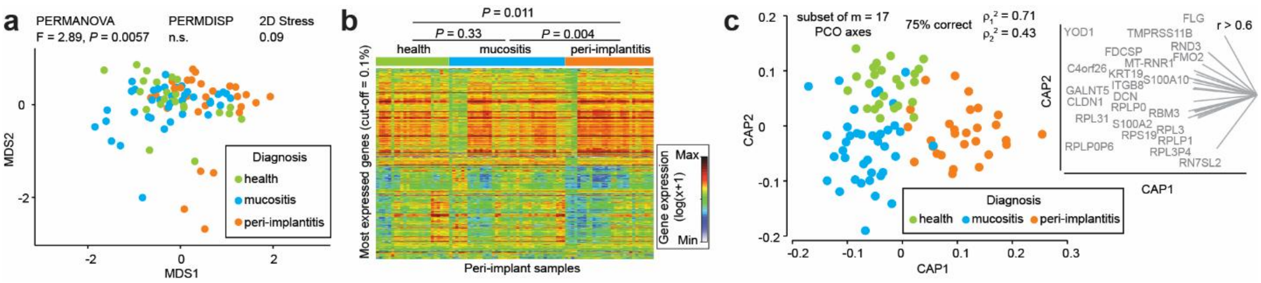


**Supplemental Figure 6. Host gene expression across diagnosis groups.** **a**, Unconstrained ordination for host gene expression data. MDS plot calculated for 91 implants on Bray-Curtis dissimilarities for log (x+1)-transformed read counts grouped to highly expressed host genes (relative activity cut-off = 0.1%). **b**, Heatmap for highly expressed genes. First, samples and genes were each clustered, and next samples were sorted by diagnosis. **c**, Diagnosis-constrained ordination for host gene expression data. CAP plot calculated on same input data as for “a”. Selected diagnosis-correlated genes (∣r∣ > 0.6) in CAP are indicated as vectors.

### Supplemental data files

**Supplemental Table 2.** Overview of full-length 16S rRNA gene reads and RNAseq reads per sample.

**Supplemental Table 3.** Class-level taxonomic profiles across diagnosis groups. Relative abundances of genera, statistical comparisons between clinical groups (Kruskal-Wallis P-values), and LDA effect sizes from CAP analysis.

**Supplemental Table 4.** Genus-level taxonomic profiles across diagnosis groups. Relative abundances of genera, statistical comparisons between clinical groups (Kruskal-Wallis P-values), and LDA effect sizes from CAP analysis.

**Supplemental Table 5.** Species-level taxonomic profiles across diagnosis groups. Relative abundances of genera, statistical comparisons between clinical groups (Kruskal-Wallis P-values), and LDA effect sizes from CAP analysis.

**Supplemental Table 6.** Differential expression of metatranscriptomic ECs across diagnosis groups.

**Supplemental Table 7.** Differential expression of ECs across microbial community types (CTs).

**Supplemental Table 8.** Average relative expression levels of functional ECs stratified by contributing taxa.

**Supplemental Table 9.** Ecological roles and metabolic associations of top 60 highly expressed ECs across diagnosis groups and CTs.

**Supplemental Table 10.** Differential expression of phage-derived transcripts across diagnosis groups.

**Supplemental Table 11.** Associations identified by HAllA (Hierarchical All-against-All) analysis between taxonomic species (16S data) and functional EC numbers (MTX data).

**Supplemental Table 12.** Associations identified by HAllA (Hierarchical All-against-All) analysis between taxonomic species (16S data) and phage transcripts profiles (MTX data).

**Supplemental Table 13.** Associations identified by HAllA (Hierarchical All-against-All) analysis between taxonomic species (16S data) and host genes transcripts (MTX data).

**Supplemental Table 14.** Associations identified by HAllA (Hierarchical All-against-All) analysis between functional EC numbers (MTX data) and host genes transcripts (MTX data).
